# An optical brain-machine interface reveals a causal role of posterior parietal cortex in goal-directed navigation

**DOI:** 10.1101/2024.11.29.626034

**Authors:** Ethan Sorrell, Daniel E. Wilson, Michael E. Rule, Helen Yang, Fulvio Forni, Christopher D. Harvey, Timothy O’Leary

## Abstract

Cortical circuits contain diverse sensory, motor, and cognitive signals, and form densely recurrent networks. This creates challenges for identifying causal relationships between neural populations and behavior. We developed a calcium imaging-based brain-machine interface (BMI) to study the role of posterior parietal cortex (PPC) in controlling navigation in virtual reality. By training a decoder to estimate navigational heading and velocity from PPC activity during virtual navigation, we discovered that mice could immediately navigate toward goal locations when control was switched to BMI. No learning or adaptation was observed during BMI, indicating that naturally occurring PPC activity patterns are sufficient to drive navigational trajectories in real time. During successful BMI trials, decoded trajectories decoupled from the mouse’s physical movements, suggesting that PPC activity relates to intended trajectories. Our work demonstrates a role for PPC in navigation and offers a BMI approach for investigating causal links between neural activity and behavior.

## Introduction

During behavior, the brain operates in a continual feedback loop with the external environment. For example, in navigation, neural activity representing planned trajectories results in physical movement, leading to updated sensory signals, which in turn inform subsequent actions. The closed-loop nature of neural activity and behavior means it is difficult to establish by measurement alone whether a signal in the brain drives a behavioral outcome, or simply reports it. This is particularly challenging in brain areas that are distant from the periphery, where neural activity may not directly relate to raw sensory or motor signals.

Previous work has identified compass- and map-like representations in multiple brain circuits^1–3^ that influence goal-directed behavior. Across species, experimental evidence implicates posterior parietal cortex (PPC) in moment-to-moment planning and execution of navigational trajectories^4–13^. Optogenetic and pharmacological inhibition of PPC impairs performance on navigation-based decision tasks without major effects on sensory processing or movement^5,14–16^. In addition, PPC has cell type-specific signals for course corrections during navigational errors^17^.

Consistent with a hypothesized role in coordinating planned movement with respect to external cues, PPC neurons carry a wide range of signals related to visual cues, environmental location, heading direction, locomotor movements, and behavioral choices^15,18–24^. Anatomically, PPC sits amidst a densely interconnected network of cortical and subcortical areas that each contribute to decision-making and navigation^19,25–29^. Together, these factors have made it challenging to pinpoint causal roles for PPC in navigation.

A strong test of whether naturally occurring activity patterns in PPC can drive task-relevant behavior is to couple these activity patterns directly to behavioral actions in real time, shortcutting pathways to motor output. In this work, we implemented this coupling using a brain-machine interface (BMI) that preserves the mapping between PPC activity and behavioral actions during navigation. We switched between navigation controlled by the mouse’s physical movements and navigation driven by BMI. We reasoned that this switching probes whether a neural signal causes a particular action at any given moment and enables us to test whether naturally occurring activity patterns are sufficient for task completion.

BMI studies often exploit the flexibility of the brain in adapting to new means of interacting with the world. Circuits that are partially involved in a behavior can adapt, and eventually control behavior proficiently, even if the BMI distorts the mapping between neural activity and behavior. For these reasons, extensive behavioral training periods are utilized, allowing neural plasticity to improve control under BMI^30–34^. We explicitly avoided these training periods and instead alternate directly between BMI and physical control, thus asking whether naturally occurring activity can drive behavioral output in the absence of learning or adaptation to the BMI.

We developed a calcium imaging-based BMI in mice to directly couple PPC activity to movement in a learned virtual reality navigation task. We found that running velocity and virtual heading direction can be decoded from PPC activity, and that these signals enabled mice to navigate with BMI control. Successful task completion was observed from the onset of switching to BMI control, without any obvious learning phase. On BMI trials, mice made apparent corrective running movements during inferred deviations in their trajectories. A significant component of the neural activity for BMI control was unrelated to these running movements, indicating that PPC has separable components of population activity for navigational running and orienting toward a goal. Our results demonstrate a causal role for PPC activity in driving navigational behavior in real time, complimenting existing work that demonstrates its necessity through silencing or lesioning^35,36^.

## Results

### Decoding PPC activity for BMI-based navigation

We trained mice to perform a decision task by navigating a virtual reality T-maze (corridor length: 225 cm) using a spherical treadmill^5^ (**Figure 1a**). During each trial, mice ran down a virtual corridor (mean time, 13.5 ± 1.4 seconds) and were rewarded for a left or right turn associated, respectively, with a black or white visual cue on the wall at the start of the maze. After weeks of training, mice learned to perform the task with high accuracy and completed hundreds of trials per session.

**Figure 1:**
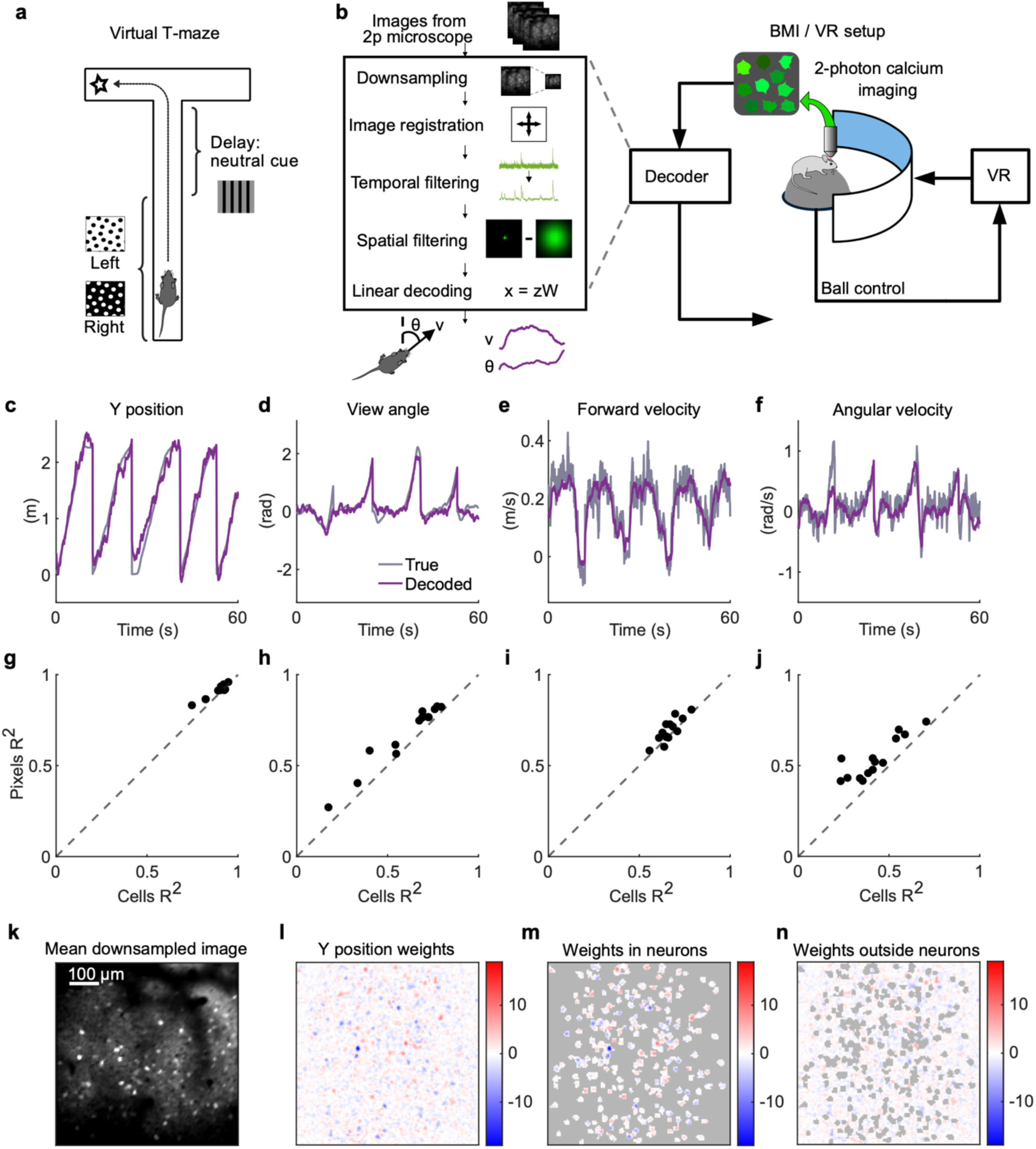
Decoding behavioral variables from neural populations in PPC. **a)** Diagram of the T-maze task. **b)** Schematic of neural and behavioral recording rig, and outline of neural data preprocessing and decoding. **c-f)** Example offline decoding of a range of variables for mouse 4. Example true (grey) and decoded (purple) values for y position, view angle, forward velocity and angular velocity are shown. **g-j)** Summary of offline decoding performance for the same variables in **c-f**, comparing decoding performance for pixels versus cells on held-out test data; data points are single sessions. **k)** example FOV after downsampling. **l)** Example of the Y position weights for the FOV in **(k)**. **m)** Same as **(l)** but only the weights inside segmented neurons. **n)** Same as **(l)**, but only the weights outside segmented neurons.

During the task, we measured activity from populations of neurons in layer 2/3 of PPC using two-photon calcium imaging of the genetically encoded calcium indicator jGCaMP7f^37^ (**Figure 1b**). To establish whether PPC activity might be suitable for use as a control signal for a virtual-navigation BMI, we first tested whether virtual heading direction and forward running speed could be decoded accurately in real time, using data from trials in trained mice. We aimed for an approach that could use data from a small number of trials, obtainable in a single session, and that could decode at a latency no greater than the imaging frame rate (30 Hz).

Our approach consisted of a linear decoder applied to imaging frames following several image processing steps (details in Methods). Briefly, we motion-corrected each image frame and down-sampled image resolution by a factor of four. Next, we temporally filtered pixel intensity to suppress changes in baseline fluorescence and high frequency noise, normalizing signals as relative changes in fluorescence (ΔF/F). We applied a spatial bandpass filter to each frame to isolate sources of calcium transients with a spatial frequency corresponding to cell bodies and to remove low spatial frequency components, such as background fluorescence. On the resulting image frames, we trained separate linear regression models for virtual heading direction, running velocity, and maze position, using iterative least mean squares (LMS).

We assessed decoder performance by comparing measured virtual trajectories in the maze to decoder predictions using held-out test data. Across mice and sessions, decoders accurately predicted maze position, virtual heading direction, forward velocity, and angular velocity (**Figure 1c-f**). Decoding performance was consistently better for positional variables than for velocity variables (maze position: R^2^ = 0.91 ± 0.01 and virtual heading direction: R^2^ = 0.69 ± 0.06; forward velocity: R^2^ = 0.67 ± 0.03 and angular velocity: R^2^ = 0.53 ± 0.04; *p*_boot_ < 0.001 for forward position vs. forward velocity; *p*_boot_ = 0.015 for virtual heading direction vs. angular velocity; N = 14 sessions from 5 mice). Decoding latency allowed us to operate at 30 imaging frames per second.

We used pixelwise intensity across the full image frame as the signal for decoding to avoid processing speed and accuracy issues that can result from automated segmentation into putative cells. Nevertheless, we compared performance of decoding using full image frames versus putative cell body regions-of-interest (ROIs) automatically segmented with Suite2p^38^ (**Figure 1g-j**). Decoding with the full image frame provided slightly better performance than with ROIs (full frame: R^2^ = 0.91 ± 0.01, 0.69 ± 0.06, 0.67 ± 0.03, 0.53 ± 0.04 for forward position, virtual heading direction, forward velocity, angular velocity vs. ROIs: R^2^ = 0.89 ± 0.02, 0.62 ± 0.06, 0.65 ± 0.02, 0.41 ± 0.04 for cells; *p*_boot_ < 0.001, < 0.001, = 0.007, < 0.001, N = 14 sessions from 5 mice). For full-frame decoding, large weights were assigned both to pixels within and outside identified neuron ROIs (**Figure 1k-n**). Thus, full-frame decoding likely extracts signals from axons and dendrites, consistent with previous work showing that signals in neuropil are predictive of behavioral state^39^.

To implement closed-loop control, we chose forward running speed and virtual heading direction as control signals. Maze position and turning velocity presented challenges that made them less suitable. Although position was decoded accurately, the decoded values showed discontinuities across adjacent time steps, causing jerky virtual motion and making it less suitable as a control signal (**Figure S1a**). We did not use turning velocity because accumulated decoding errors in angular velocity could be problematic, as mice must maintain a narrow range of heading angles to successfully run down the stem of the maze (**Figure S1b**).

### Mice reliably navigate to goal locations in virtual environments using the BMI

We integrated our decoder into the microscope acquisition software^40^ to enable switching between ball and BMI control during experiments. In “Ball trials”, rotation of the treadmill updated motion in the virtual environment. In BMI trials, the forward velocity and virtual heading direction decoded from neural activity were routed to the virtual reality engine, bypassing the treadmill signal (**Figure 2a**).

**Figure 2:**
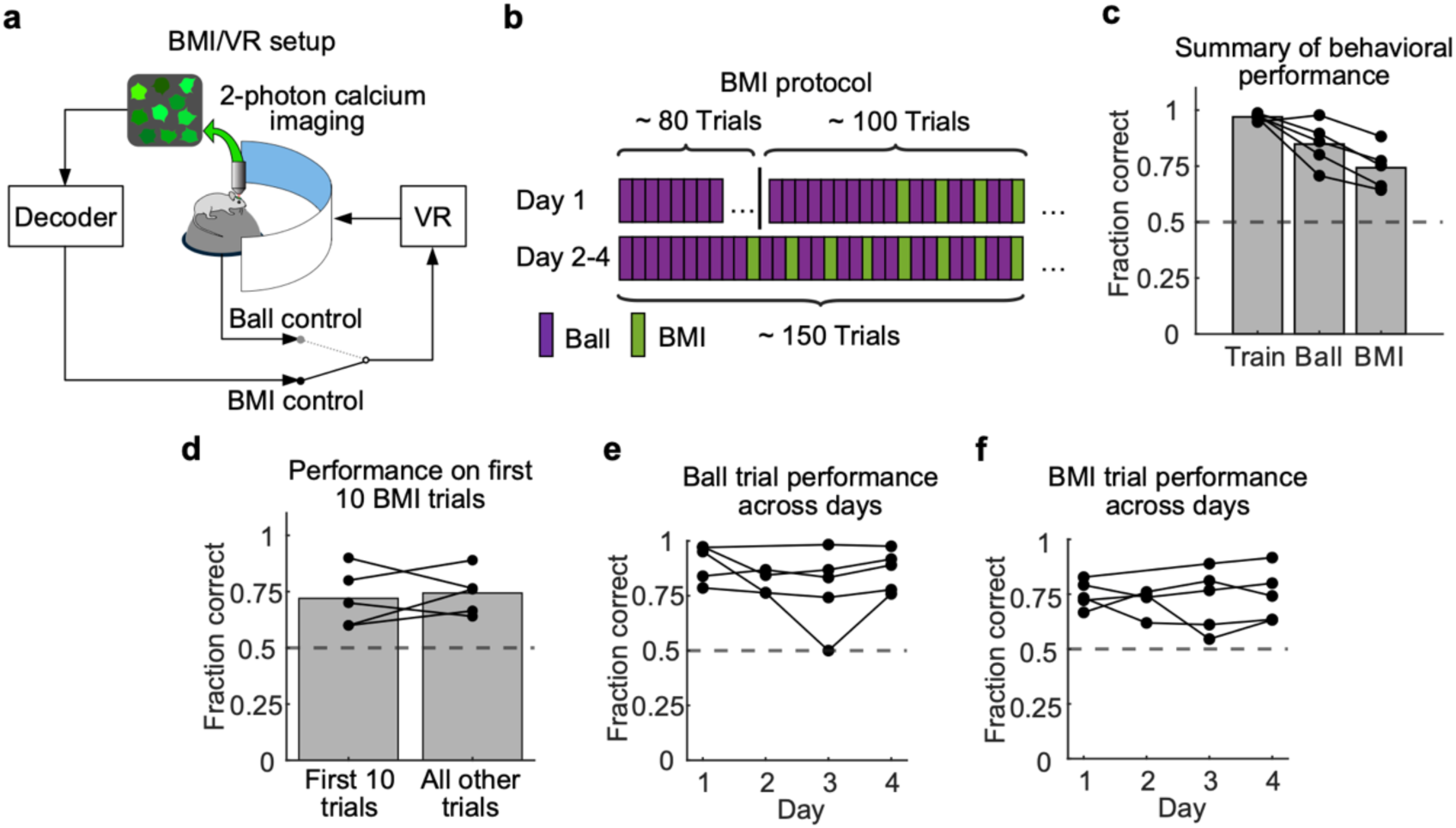
Mice perform trials of the virtual navigation task using an optical BMI. **a)** Schematic of closed-loop BMI experiment setup **b)** Outline of trial structure during BMI training and testing. **c)** Summary of task performance during training and testing sessions, measured as the fraction of correctly completed trials. Lines are individual mice, bars are means across mice. **d)** Performance on first ten versus all subsequent BMI trials. **e)** Task performance of mice on ball trials across days of experiments. Lines are individual mice. **f)** Task performance of mice on BMI trials across days of experiments. Lines are individual mice.

Mice were trained to expert level (>90% correct trials) before commencing sessions with BMI trials. Sessions with BMI trials began with a block of trials in which the mouse navigated the virtual maze using the treadmill (∼80 trials, **Figure 2b**). We used imaging data acquired during this period to train the BMI decoder. In subsequent trials, we interleaved BMI trials with Ball trials, with two consecutive Ball trials for every BMI trial (**Figure 2b**). We analyzed BMI performance from the first trial in which BMI control was introduced, without any specific cue or behavioral training for BMI use. After training the decoder, we fixed the decoding weights for the remainder of the experiment, including for sessions on subsequent days.

Mice were able to complete BMI trials without major aberrations in trajectories through the maze and at similar rates as for Ball trials. Behavioral performance in BMI trials approached levels observed in Ball trials (**Figure 2c**). Remarkably, performance was above chance from the onset of BMI use in the first session (p = 0.0013 using a binomial test for the first 10 trials from all 5 mice). Mice that were naïve to BMI performed with similar accuracy during the first 10 BMI trials as they did in subsequent BMI trials (**Figure 2d, Figure S2a**), indicating that PPC activity drove navigational behavior in BMI without any obvious learning. Although BMI performance was worse in some mice, when combining trials from all BMI testing sessions for each mouse, all mice performed significantly above chance on BMI trials (p = 3.14×10^-13^, 9.21×10^-6^, 1.72×10^-4^, 0, 9.51×10^-12^, for all mice using a binomial test with N = 198, 180, 160, 119, 146 BMI trials). We noticed that Ball trial performance was slightly lower than in trials used for training the decoder, perhaps due to mice sensing differences in the task during interleaved BMI trials.

Previous work reported that tuning to behavioral variables for individual neurons changes over time in mouse PPC^15,20^. Such changes could lead to accumulating decoding error in BMI. However, during the five days spanned by our experiments, behavioral performance on BMI and interleaved Ball trials remained constant (**Figure 2e-f, Figure S2c-d**).

Taken together, these results show that mice can successfully navigate to goal locations using activity decoded from PPC in real time, and without a period for learning. Thus, naturally occurring PPC activity patterns are sufficient to drive navigational heading in this experimental setting. This result is not a trivial consequence of the observations in Figure 1, or of previous studies establishing correlations between navigational trajectories and neural activity in PPC.

First, accurate trajectory decoding from offline data does not guarantee successful closed-loop BMI performance. Decoding errors can accumulate in a closed-loop setting, leading to destabilization of the BMI and loss of accurate control^41^. Furthermore, decoding is imperfect, and mice may react to unanticipated deviations in the trajectory during BMI. This may in turn generate PPC activity patterns that differ substantially from the data used to train the BMI decoder, further decreasing control accuracy.

Second, in many BMI approaches, neural plasticity allows a subject to gradually learn how to control a BMI in closed loop, even if the mapping between neural activity and BMI output differs substantially from open loop calibration. By contrast, we observed that naturally occurring PPC activity patterns measured in open loop were sufficient to permit real-time decoding in closed loop with no learning phase, as we tested behavioral performance immediately upon switching to BMI control.

Third, it could have been the case that PPC activity is simply a readout of task states like maze position and heading direction, with no direct role in driving changes in navigational state. We believe this is unlikely, because updating a trajectory in real-time requires a signal with a predictive component that would not be present if the signal simply reports the current state of the animal in the VR task. To illustrate this, we built a simple neural network model to test whether a population of units tuned for heading direction (**Figure S3a**) allow an agent to solve the T-maze task. In the first scenario, units report the agent’s current heading direction, and thus it is possible to decode heading direction from these units as the agent navigates (**Figure S3b, top**). However, when decoding the signal from such units to mimic the BMI scenario, the agent was unable to update its heading and unable to complete trials (**Figure S3b, bottom**). In an alternative scenario, we modified the model so that population activity reflects the agent’s intended heading direction in the subsequent timestep. In this case, as before, we could accurately decode the agent’s heading direction (**Figure S3c, top**). However, unlike the previous case, closed-loop decoding of this population activity enables the agent to navigate and perform the task (**Figure S3c, bottom**). Therefore, predictive signals (i.e., signals pertaining to intended actions) are essential for closed-loop BMI control. We note that predictive signals could be learned during adaptation to a BMI; however, as noted in the previous point, our data do not support such learning.

### Running trajectories differ between Ball and BMI trials

During successful BMI trials, mice traversed the T-maze and turned at the end of the maze into the rewarded arm (**Figure 3a**). The navigational trajectories through the maze on BMI trials, as dictated by the decoder output, often resembled trajectories during Ball trials, in which the mouse’s running specified the trajectory (**Figure 3b-c**). Trial duration during correct trials was comparable between Ball and BMI trials (16.5 ± 0.8 s, 15.8 ± 0.7 s, *p*_boot_ = 0.29. N = 19 sessions, 5 mice; **Figure 3d**). These observations, in addition to above-chance left-right choices at the T-intersection in BMI trials, means it is unlikely that progress through the maze during BMI resulted from random motion generated by the decoder.

**Figure 3:**
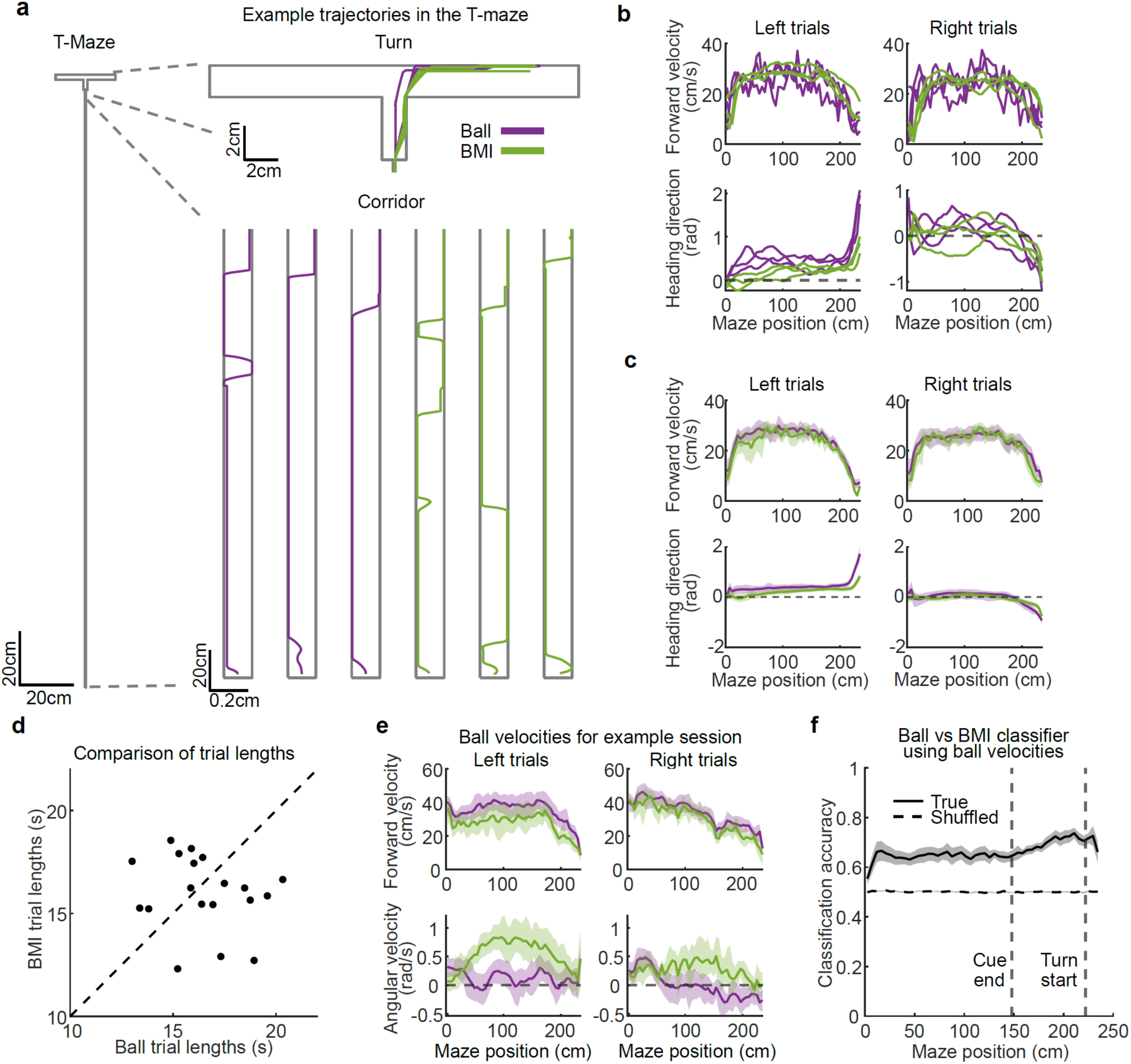
Mice generate different running trajectories on Ball and BMI trials. **a)** Example trajectories through the T-maze during ball and BMI right trials. Trajectories are shown separately in the straight corridor, and for the turn. **b)** Example forward velocity and view angle trajectories on Ball and BMI trials; For Ball trials, these values are generated by the spherical treadmill. For BMI trials, these values are generated by the decoders. **c)** Mean forward velocity and view angle trajectories for an example session, with 95% bootstrapped confidence intervals. **d)** Trial lengths for ball and BMI trials. Each point is an individual session. **e)** Mean forward velocity and angular velocity trajectories as measured on the ball for ball and BMI trials, with 95% bootstrapped confidence intervals. **f)** Classification accuracy of trials as ball or BMI using measured ball velocities. Separate SVM classifiers were trained in each spatial bin. Hierarchical bootstrap means and standard deviations across all sessions are shown for the true data (solid line) and for data with shuffled trial type labels (dashed line).

Our experimental setup allowed us to measure the mouse’s running in both Ball and BMI trials even though, by design, ball rotations had no effect on movement in the virtual maze during BMI. We first asked whether mice simply memorized stereotyped running patterns from trial to trial, in which case the running trajectories are expected to be highly similar between Ball and BMI trials. In this case, the mouse could generate stereotyped neural activity sequences, leading to successful completion of BMI trials, irrespective of interaction with the virtual environment. Such “autopilot” behavior could confound the interpretation that PPC activity controls immediate actions in BMI trials because in this case PPC activity might instead report stereotyped motor outputs. Multiple lines of evidence argue against this “autopilot” possibility.

During BMI trials, all mice were able to generate course corrections when they began to turn in the wrong direction in the T-stem (Examples, **Figure S4**), indicating real-time adjustments in course instead of only stereotyped trajectories. Also, apart from a small minority of trials (**Figure S5a**), running trajectories were detectably different in Ball and BMI trials (e.g. **Figure S5b**). Yaw treadmill velocities, which control virtual heading direction in Ball trials, differed substantially (e.g., **Figure 3e**; change in correlation across trial types: −0.041 ± 0.007 for forward velocity, −0.10 ± 0.01 for angular velocity, *p*_boot_ <.001; N = 19 sessions from 5 mice; **Figure S5c**). In some BMI trials, mice generated large, continuous yaw rotations on the treadmill throughout the trial (e.g. **Figure S5d**). These ball rotations would generate unidirectional rotation of the virtual heading and prevent progress through the maze. Nevertheless, through BMI control, the neural activity in PPC in these trials generated normal trajectories toward the goal location. Forward treadmill velocity profiles tended to be more similar between Ball and BMI trials, as expected given that the task proceeds by running without pause toward a goal.

To further compare running trajectories on Ball and BMI trials, we trained a support vector machine (SVM) classifier to predict whether a given trial was a Ball or BMI trial from the running velocities alone. We trained separate classifiers for left-choice and right-choice trials and excluded incorrect trials. At all spatial bins after trial onset, the classifier distinguished Ball and BMI trials using the instantaneous rotational velocities of the treadmill (All bins significant after FDR correction at α = 0.05; N = 19 sessions from 5 mice; **Figure 3f**). Together, these results indicate that mice do not repeat stereotyped running motions from one trial to the next and instead steer their trajectories in real-time during BMI. Further, the dissociation between the mouse’s physical movements and the decoder’s predicted heading direction suggest that the decoder relies on neural activity related to intended heading direction rather than activity for running movements.

### Mice attempt corrective running for heading deviations during BMI trials

Monitoring running on the ball during BMI trials enabled us to infer reactions to BMI output. We reasoned that if BMI output deviates from a mouse’s expectations, we may detect reactive movements through changes in the treadmill velocity. To infer deviations in expected trajectories, we compared BMI trajectories with the mean trajectories gathered under normal (Ball) conditions.

We focused on deviations in heading direction because these are more easily perceptible and task-critical than variations in forward velocity. In each spatial bin of correct BMI trials, we computed the deviations in virtual heading from the mean virtual heading in correct Ball trials^17^ (**Figure 4a**). We unavoidably included any natural trial-to-trial variation in behavior as part of this error, but reasoned that large, systematic errors would result in a detectable deviation for which the mouse might correct. Attempts to correct for these heading deviations would manifest as correlated angular velocity changes on the ball with opposite sign to the error (example, **Figure 4b**). Indeed, we found heading deviation and ball angular velocity to be negatively correlated across sessions and mice (bins 4–50 significant after FDR correction at α = 0.05; N = 19 sessions from 5 mice; **Figure 4c**).

**Figure 4:**
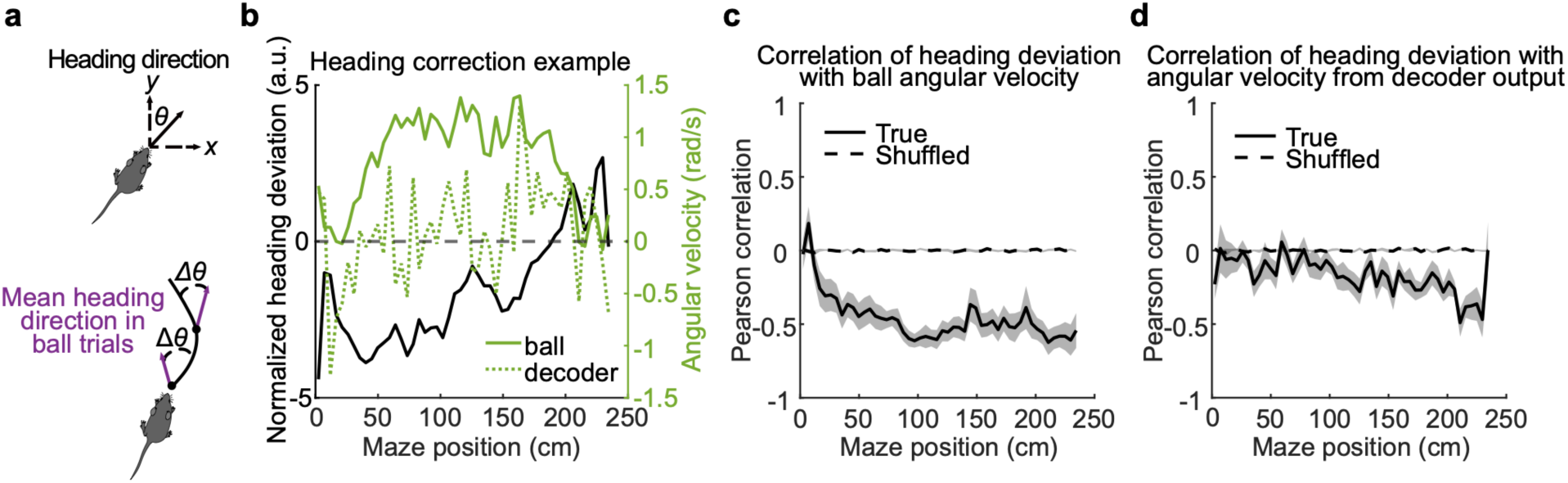
Mice attempt to use the ball to correct for heading deviations during BMI trials. **a)** Diagram of how heading deviation was estimated during BMI trials. The purple arrow indicates average heading in that spatial bin during ball trials. **b)** Example of spatially binned heading deviation, and angular velocity, either measured from the ball, or calculated from decoder output, for a single trial. **c)** Correlation between heading deviation and ball angular velocity on BMI trials for different spatial bins throughout the maze. Hierarchical bootstrap means and standard deviations are shown for the true data and for shuffled data. **d)** Correlation between heading deviation and angular velocity calculated from decoder output on BMI trials for different spatial bins throughout the maze. Hierarchical bootstrap means and standard deviations are shown for the true data and for shuffled data.

Interestingly, even though mice changed running to correct for these heading deviations, the BMI output often did not make rapid corrections to the heading direction. To estimate heading corrections in the BMI output, we differentiated the BMI heading direction signal to obtain a measure of angular velocity from the decoder output (example, **Figure 4b**). The correlation between the decoder heading changes and the heading deviation was weaker than between the mouse’s running velocity and the heading deviation (23/50 bins significant after FDR correction at α = 0.05; N = 19 sessions from 5 mice; **Figure 4d**). Despite the lack of instantaneous heading corrections by the BMI output, over the entire trial, the BMI output necessarily steered the correct overall path toward the end of the maze, because we only included correct trials in this analysis.

The mice’s attempts to correct for transient heading deviations through running indicate decoupling of immediate physical movements from the decoder output. This decoupling, combined with the fact that mice successfully perform BMI trials, suggests that the decoder acts on neural activity related to heading direction that may deviate from immediate motor actions. Furthermore, these reactive changes in motor output, combined with the general differences in running trajectories on Ball and BMI trials reported above, further indicate that mice actively control motion via the BMI rather than repeating a memorized set of actions.

### Distinct PPC activity for navigational turning and heading

The transient decoupling between BMI decoding and corrective running movements during large heading deviations raises two possibilities. Neural signals corresponding to these corrective movements are not present in PPC. Alternatively, these signals are present but distinct from the signals used by the decoder for heading direction.

To test these possibilities, we analyzed neural activity when mice made physical reactions to heading deviations in BMI trials. Some neurons had activity that was correlated with the magnitude of the ball angular velocity during BMI trials (example ρ = 0.56, *p* = 3.34×10^-16^ N = 180 binned samples; **Figure 5a**). PPC population activity also tended to increase during corrective movements (example session shown in **Figure 5b**, hierarchical bootstrap across all sessions: ρ = 0.23 ± 0.05, *p*_boot_ < 0.001, N = 19 sessions from 5 mice). Thus, PPC contains activity related to the physical movements associated with reactions to heading deviations during BMI control.

**Figure 5:**
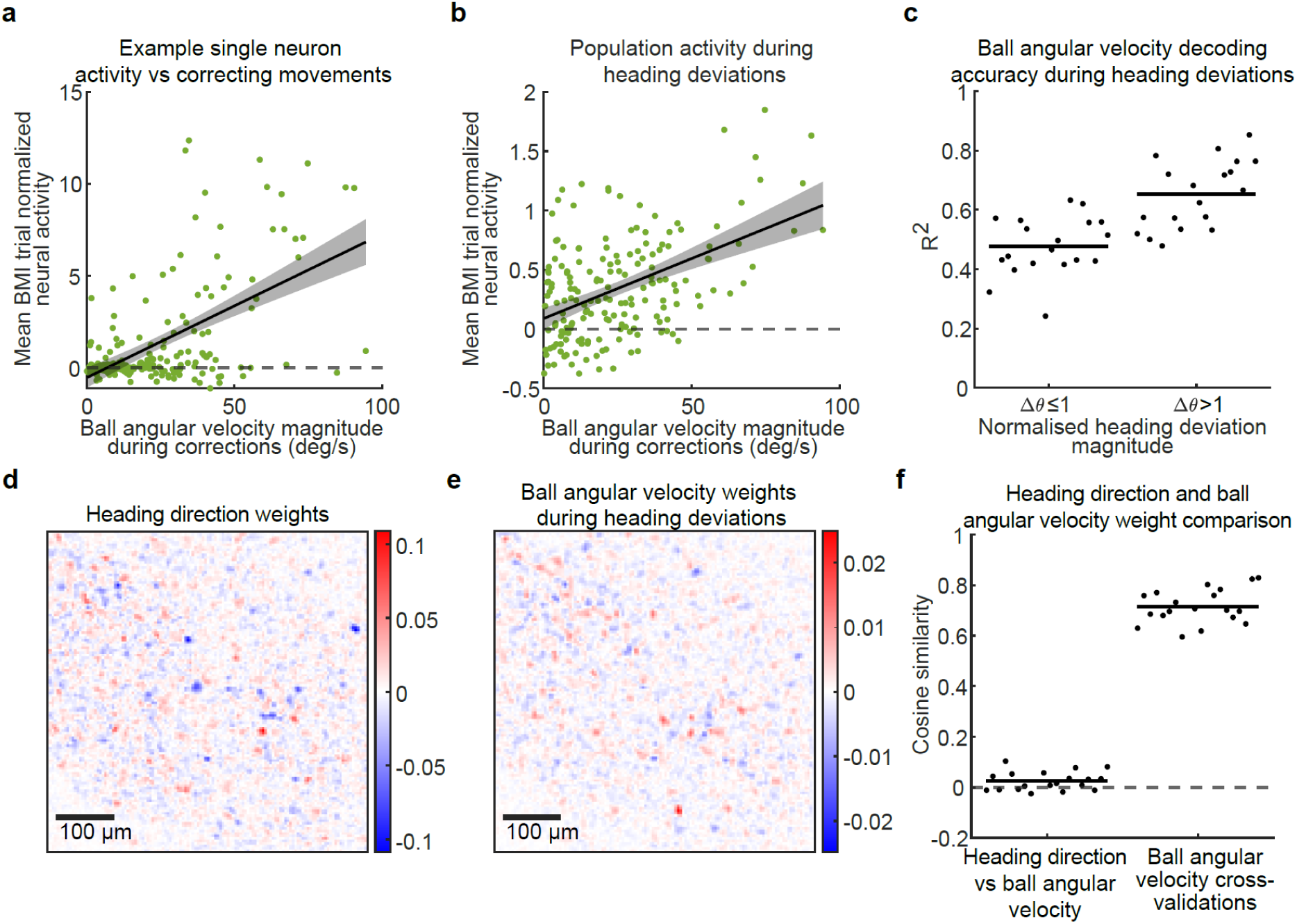
BMI-related neural activity for setting heading direction is distinct from activity patterns related to ball movements during course corrections. **a)** Mean normalized activity on BMI trials for a single neuron plotted against ball angular velocity magnitude in bins with a heading deviation and where angular velocities were heading correcting. Error bars are 95% confidence intervals. **b)** Mean normalized population activity during BMI trials plotted against ball angular velocity in bins where this is a heading deviation and where angular velocities were heading correcting, as in **a**. Error bars are 95% confidence intervals. **c)** Ball angular velocity decoding accuracy of cross-validated decoders trained on angular velocities during BMI trials and tested separately on samples with normalized heading deviations less than or equal to 1, and greater than 1. **d)** Example BMI heading direction decoding weights for one imaging plane for one mouse. **e**) Example ball angular velocity weights for one imaging plane trained offline on BMI trials for a single session for the same mouse as (**d**). **f)** Cosine similarities between BMI heading direction weights and offline ball angular velocity weights (left), with cross-validated cosine similarities of offline ball angular velocity weights (right) for comparison. Each point = single session, black lines = means.

We reasoned that the decoder may have failed to capture this aspect of PPC activity related to the reactive physical movements because it reflects a different aspect of neural activity. We trained offline decoders on neural activity from BMI trials to test whether corrective ball angular velocity movements could be reliably extracted. Angular motion that was in the direction aimed toward correcting for heading deviations could be decoded accurately, with better decoding accuracy for large deviations (small normalized heading deviations, |Δ*θ*| ≤ 1: R^2^ = 0.48 ± 0.03, larger normalized heading deviations, |Δ*θ*| > 1: R^2^ = 0.66 ± 0.04. *p*_boot_ < 0.001. N = 19 sessions from 5 mice; **Figure 5c**). We compared weights in the BMI decoder (which extracts heading direction) with this decoder trained to output angular velocity. The two decoders tended to assign their largest weights to different pixels in movies of PPC activity (example, **Figure 5d-e**). We quantified the similarity of these decoder weights by computing the cross-validated cosine similarity of the two weight vectors. The cosine similarities were low between the weights in the heading direction and angular velocity decoders (0.026 ± 0.01. N = 19 sessions from 5 mice; **Figure 5f**). As a control, decoder weights were similar across 30 different random subsamples of the ball angular velocity decoder in BMI trials (cosine similarity = 0.72 ± 0.02; N = 19 sessions from 5 mice). Thus, there is little apparent overlap between the dimensions in neural activity that are most useful for decoding heading direction and those for ball angular velocity. However, it may be possible to identify other decoders with different sets of weights and similar decoding performance that exhibit greater overlap between PPC activity for running movements and heading direction.

These analyses show that the PPC contains linearly separable signals for running-related movements and navigational signals (i.e., heading direction). Combined with the observation that BMI output deviated significantly from the mouse’s physical running movements, this suggests that in training our BMI heading direction decoder we identified a specific subspace of PPC activity that relates to navigational heading and that is largely independent of signals related to physical running movements. This explains why the navigational trajectory generated by the BMI output could substantially deviate from the mouse’s running outputs while still maintaining a correct overall path toward the goal location.

### BMI trials have elevated PPC activity levels

Behavior and task performance differed between Ball and BMI trials, suggesting that neural activity also differed between these trials. Indeed, SVM classifiers accurately distinguished Ball and BMI trials at all spatial bins in the maze using population activity (All bins significant after FDR correction at α=0.05; N = 19 sessions from 5 mice; **Figure 6a**). The difference in neural activity between Ball and BMI trials was not due to large re-tuning in neural activity. In both Ball and BMI trials, PPC activity was organized as trial type-specific sequences across BMI and Ball conditions, with relatively consistent neural tuning across the duration of the trial (**Figure 6b**).

**Figure 6:**
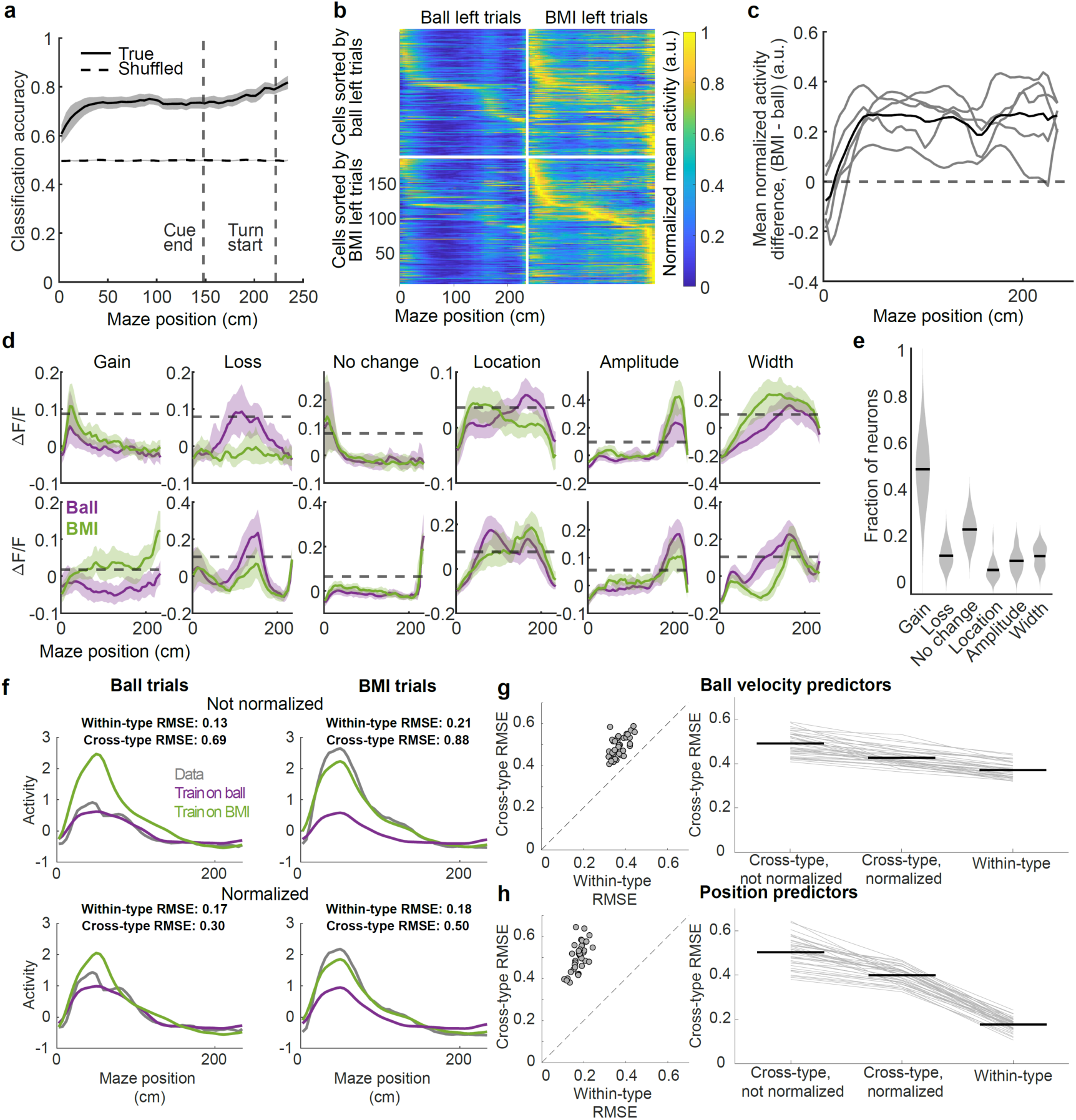
Neural activity increases in BMI trials. **a)** Ball vs BMI classification accuracy for SVM classifier on neural activity Hierarchical bootstrap means and standard deviations across all sessions are shown for the true data (solid line) and for data with shuffled trial type labels (dashed line). **b)** Normalized activity raster plots from an example session for ball (left column) and BMI left trials (right column) sorted according to the location of peak activity in ball trials (top row), or BMI trials (bottom row). Cell order is maintained across ball and BMI conditions within a row. **c)** Mean population activity difference (BMI – ball) as a function of maze location. Grey lines = means for individual mice; black line = group mean. **d)** Example neural tuning curve comparisons for BMI vs ball conditions. Bootstrapped means and 95% confidence intervals are shown. **e)** Distribution of classes of differences in neural tuning curve in BMI vs ball conditions across all animals and sessions. **f)** Example single-neuron linear model fit using position to predict neural activity. Values are RMSE. **g)** For ball velocity predictors: prediction error for models trained and tested within or across trial types (left) and mean RMSE values for cross-type, cross-type with normalization, and within-type predictions. **h)** Same as (g) but for positional predictors.

The most apparent difference was elevated activity in BMI trials relative to Ball trials, as revealed when normalizing activity levels consistently across both trial types. Elevations in mean activity in BMI trials relative to Ball trials were evident almost immediately after mice began BMI trials and persisted for the remainder of the trial (**Figure 6c**). These BMI-related increases in activity were widespread across the population (fraction of neurons with increased activity on BMI trials = 0.79 ± 0.05, N = 19 sessions from 5 mice; **Figure S6a**) and consistent across trials within a day and across days (**Figure S6b-c**). We investigated whether differences in running patterns between Ball and BMI trials explained BMI-related increases in neural activity. Neurons with larger weights in the ball angular velocity decoder tended to exhibit larger increases in activity on BMI trials, in contrast to neurons with larger weights in the heading direction decoder (**Figure S6d**).

We asked whether excess BMI activity might be due in part to the corrective movements mice perform during heading deviations (as described in **Figure 4**). We found a significant correlation between ball angular velocity and excess neural activity in BMI trials relative to Ball trials during corrective actions (*ρ* = 0.18, p = 3e-74, **Figure S7a**). Consistent with our analyses from Figure 5, this excess activity did not generate changes in heading direction in the BMI decoder (**Figure S7b,** no bins significant after FDR correction at α=0.05; N = 19 sessions from 5 mice).

Changes in tuning were modest between Ball and BMI trials, as characterized by the location in the maze at which neurons had their peak of activity. The most common change was a gain of a peak of activity during BMI trials, with fewer cells showing a loss of a peak or a change in peak location or width (**Figure 6d-e**). We further tested for tuning shifts between Ball and BMI trials by training linear models to predict the activity of individual neurons using either maze position or the pitch, roll, and yaw velocities of the ball as predictors. We used a subset of Ball or BMI trials as a training set and tested on held-out trials of the same or different type (examples, **Figure 6f**). Due to the increases in mean activity on BMI trials, we trained one set of models that maintained these activity shifts and another set in which changes in mean activity were minimized by normalizing activity within each trial type. Models performed worse when tested on the different trial type than on the same trial type (e.g., train on Ball trials/test on BMI trials vs. train on Ball trials/test; p_boot_ < 0.001 for position or ball velocity predictors, N= 19 sessions from 5 mice; **Figure 6g-h**). The models trained using position predictors had worse performance across trial types than the models trained using ball velocities, suggesting that neurons changed more in their position tuning than their velocity tuning (**Figure 6g-h**). Accounting for mean activity between Ball and BMI trials improved model performance when tested on each trial type, confirming that shifts in mean activity occur between Ball and BMI trials (**Figure 6g-h**).

Changes in population activity between BMI and Ball conditions could reflect a reorganization of the internal relationships between cells, or simply a change in the network state. To discriminate between these possibilities, we computed pairwise correlations in activity between cells, across trials, conditioned on maze position (commonly known as “noise correlations”). We found this measure to be largely preserved between Ball and BMI conditions (**Figure S6e-g**), supporting a change in network state, as opposed to a BMI-dependent reorganization of relationships between cells.

## Discussion

Our findings contribute to a growing understanding of how navigational plans are represented in the cortex, by demonstrating that a mouse’s intended trajectory can be read out from PPC activity, and that these predictive signals can steer complex behavior in real-time via a BMI. Previous work revealed a complex mix of sensory, motor, and cognitive variables encoded in PPC, as well as other cortical regions^19,25,26^. Such complexity is expected because goal-oriented actions necessarily depend on kinematic, sensory, and context-dependent variables, and all of these depend on an animal’s internal state as well as the state of the environment. In the PPC, neural signals correlate with heading, position, and intended path with respect to an external goal, permitting offline reconstruction of behavioral trajectories from neural recordings^15,20,22,25,26,42^. Furthermore, inactivation and disruption of activity in PPC indicate this area is necessary for mice to perform navigation and decision-making tasks^5,14,43^.

Together with previous work, our findings support a general theory that neural activity in PPC contributes to high-level planned trajectories, particularly those associated with navigating toward a goal^10,12,26^. Specifically, we demonstrated that real-time navigation can be driven by directly coupling movement in a VR environment to predictive signals extracted from PPC. This coupling bypasses pathways that would normally transform intended trajectories into movement through cortical, subcortical, and spinal pathways. This provides a strong test of whether a fixed, inferred mapping between PPC activity and executed actions remains intact under direct coupling, and whether PPC activity is potentially sufficient for controlling behavior.

There were many reasons to expect that mice may fail in the BMI version of the virtual task, especially during initial exposure to BMI control. The task is complex, requiring correct decisions and extended trajectories over more than ten seconds and hundreds of centimeters. Under normal, non-BMI conditions, mice require four to six weeks of daily training for performance to plateau. Despite this, mice managed to immediately navigate using the BMI, without any observed requirement for learning to use the BMI to control virtual navigation. We interpret this as evidence that the PPC signals we used to train the BMI decoder do indeed represent planned actions on a moment-to-moment basis.

From neural recordings alone, it remained a possibility that activity in PPC passively reports the current or recent environmental and behavioral state. Under this interpretation, PPC activity in learned, stereotyped tasks over many repetitions would inevitably correlate with task-relevant variables, including the mouse’s trajectory. However, such signals would have no direct role in driving actions and would not correspond to a planned trajectory. Importantly, we demonstrated in a simple model that passive readout of the behavioral and environmental state cannot provide effective real-time control in a closed-loop setting where no learning is involved. Combined with observed decoupling of physical running from BMI trajectories, this indicates that the BMI decoder did not merely extract a correlate of raw sensory or motor signals.

Despite successful task performance in the majority of trials, mice encountered greater difficulty executing the task under BMI control. There are several potentials reasons for this, and all of them reinforce the non-trivial nature of successful real-time BMI control. First, decoder predictions contain errors. In a closed-loop system, feedback means that errors easily accumulate and can lead to loss of stability of BMI control. Thus, there are complicated tradeoffs for different combinations of experimenter-set parameters, such as decoder gain and smoothing time constant. These issues are well documented in BMI studies that control kinematics of robotic arms and computer pointers^41^. In such cases, successful BMI control relies on a subject’s ability to learn and incrementally improve performance over time, often with verbal coaching. In our experiments, mice did not require time to adjust to BMI. Although task completion was poorer in BMI conditions, mice kept control of virtual motion during most trajectories, adjusting their courses appropriately in real time to complete trials successfully.

Second, we considered it a possibility that, without extensive overt learning phases, mice would disengage from the task on the interleaved BMI trials as soon as they recognized a mismatch between running on the treadmill and motion in the virtual environment. We presented BMI trials at a fixed interval of every third trial, so that mice would not receive long blocks of BMI trials. Mice performed BMI trials with consistent levels of performance (**Fig S2c-d**) within and across sessions, even from the first group of BMI trials.

Third, in many brain areas, neural population activity can remap in different behavioral contexts^44^. In PPC, different physical contexts recruit largely different sets of neurons in tasks with similar cognitive demands^45^. In our experiments, the abrupt change of control from Ball to BMI trials would likely have been detectable due to some amount of mismatch between running and motion of the virtual environment. This could have caused a contextual remapping of population activity, potentially destroying decoder performance. Notably, neural activity generally increased in PPC during BMI trials, potentially due to arousal, or detection of the BMI context. However, these changes were sufficiently compatible with the linear readout of the BMI decoder to enable controlled virtual navigation.

Our experimental setup allowed us to infer when mice attempted physical corrections for heading deviations during BMI trials. Because treadmill motion deviated from BMI output during these events, we were able to detect distinct neural activity patterns for running movements and heading direction. The corrective actions and their associated neural activity are consistent with our observation of navigational course correction signals in PPC^17^. Any inference of intended behavior is imperfect because it relies on extrapolation of typical behavioral patterns to specific episodes. However, since mice also successfully completed BMI trials even while making counterproductive movements with the ball, it is likely that the decoder extracted signals representing general heading along an intended trajectory, as opposed to immediate reactions.

Finally, previous studies found that fixed, offline decoding of behavior from PPC activity gradually degrades over the course of several days due to representational drift^15,20^. Consistent with these findings, we observed increases in offline decoder error over the four consecutive days of experiments (**Figure S8a-b**). However, mice maintained stable task performance with a fixed BMI decoder during this same period. This may be due to some amount of plasticity within PPC or due to behavioral compensation during BMI use. However, given previous observations of the rate of representational drift in PPC, a key question for future work is whether yoking activity to behavior via a BMI stabilizes representations over longer periods.

### Limitations

There are limitations in the design of our study and the data we gathered. Although mice performed above chance on BMI trials, they exhibited better performance on Ball trials. This difference could result from imperfect decoding of intended heading direction or perceptible lags in decoder output due to relatively slow calcium signals.

We chose to use a linear decoder, which is efficient and relatively interpretable. However, it is possible that more complex decoders, such as multilayer artificial neural networks, might generate a more accurate control signal from population activity. Even simple additions to a linear decoder, such as using a linear dynamical system instead of a static mapping, may improve decoder accuracy. Both alternatives would require decoding accuracy to be traded against computational speed.

Future BMI approaches may benefit from larger recorded neural populations and faster feedback between recording and behavior. We operated our BMI at the acquisition rate of the two-photon microscope (30 Hz). This rate was not necessarily a limiting factor for the BMI because physical movements on Ball trials are also subject to inertia of the treadmill. Moreover, the decay time of the calcium indicator, on the order of 100 ms, exceeds the sampling rate, placing a limit on the response of the decoder. Lower latency readout of neural signals may therefore improve BMI performance beyond what we observed.

There are limitations to our interpretation of the results. One of our goals was to fill a missing link in the causal chain between PPC activity and goal-directed navigation. Earlier work established correlations between the two and provided evidence that disruption of PPC function impacts behavior. Obtaining evidence for the converse – that PPC is in some sense sufficient – is challenging. Any experimental manipulation changes the context between PPC activity and behavior, meaning that in a strict sense one cannot draw conclusions about what “would have happened” under normal, non-experimental conditions. Thus, our inference about the sufficiency of PPC activity for controlling behavior must be interpreted in this context. We cannot conclude that PPC controls behavior in precisely the same way during BMI as it does in control conditions. However, the fact that mice completed the task far above chance with no apparent learning has a parsimonious explanation: the activity patterns we observed in control conditions map directly to actions, and control behavior. Further work along similar lines can reduce uncertainty in this conclusion.

## Supplemental Figures

**Figure S1:**
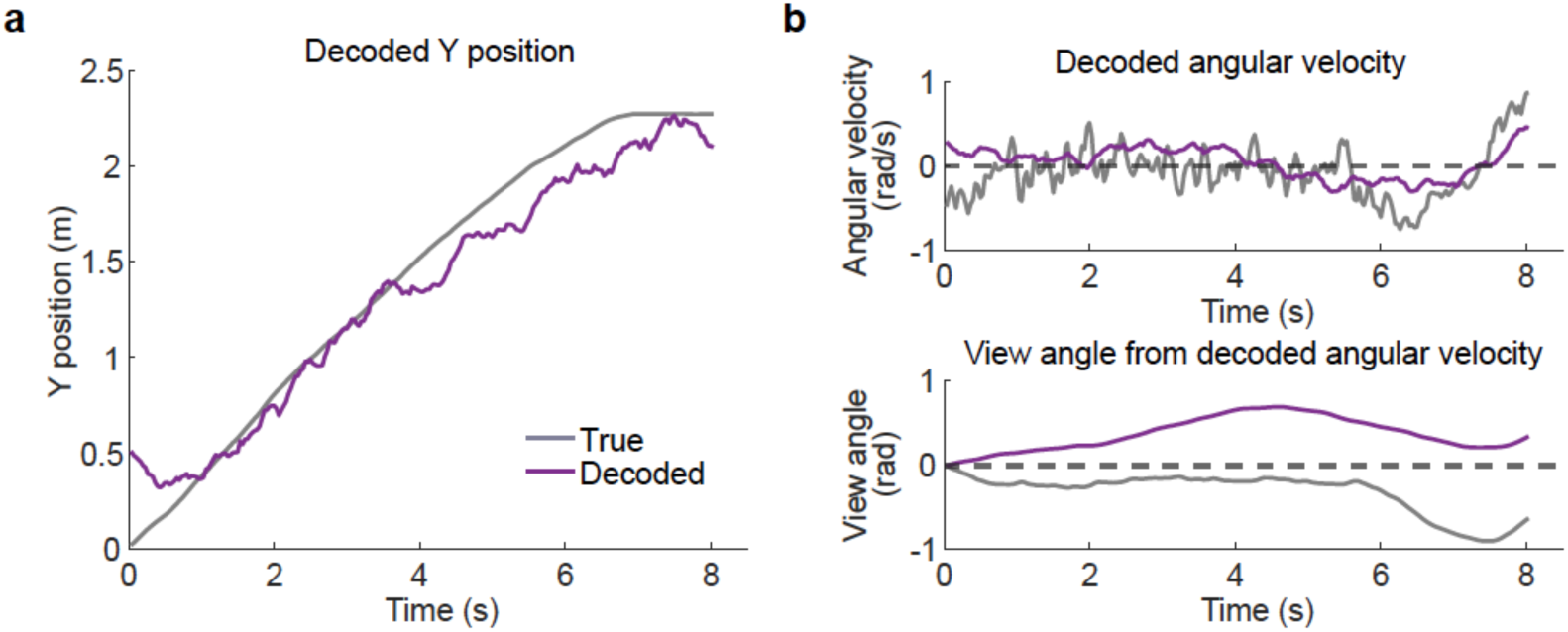
Example results from alternative control signals. **a)** Example decoding of Y position compared to true Y position for a single trial. The decoded position trajectory is discontinuous and would be difficult to use as a control signal. **b)** Example of decoded angular velocity compared to true angular velocity during a ball trial (top). Comparison of the result of integrating decoded or true angular velocity throughout the same trial (bottom). Large errors in view angle accumulate over a trial when integrating this decoded angular velocity, even when angular velocity itself is decoded with high accuracy.

**Figure S2:**
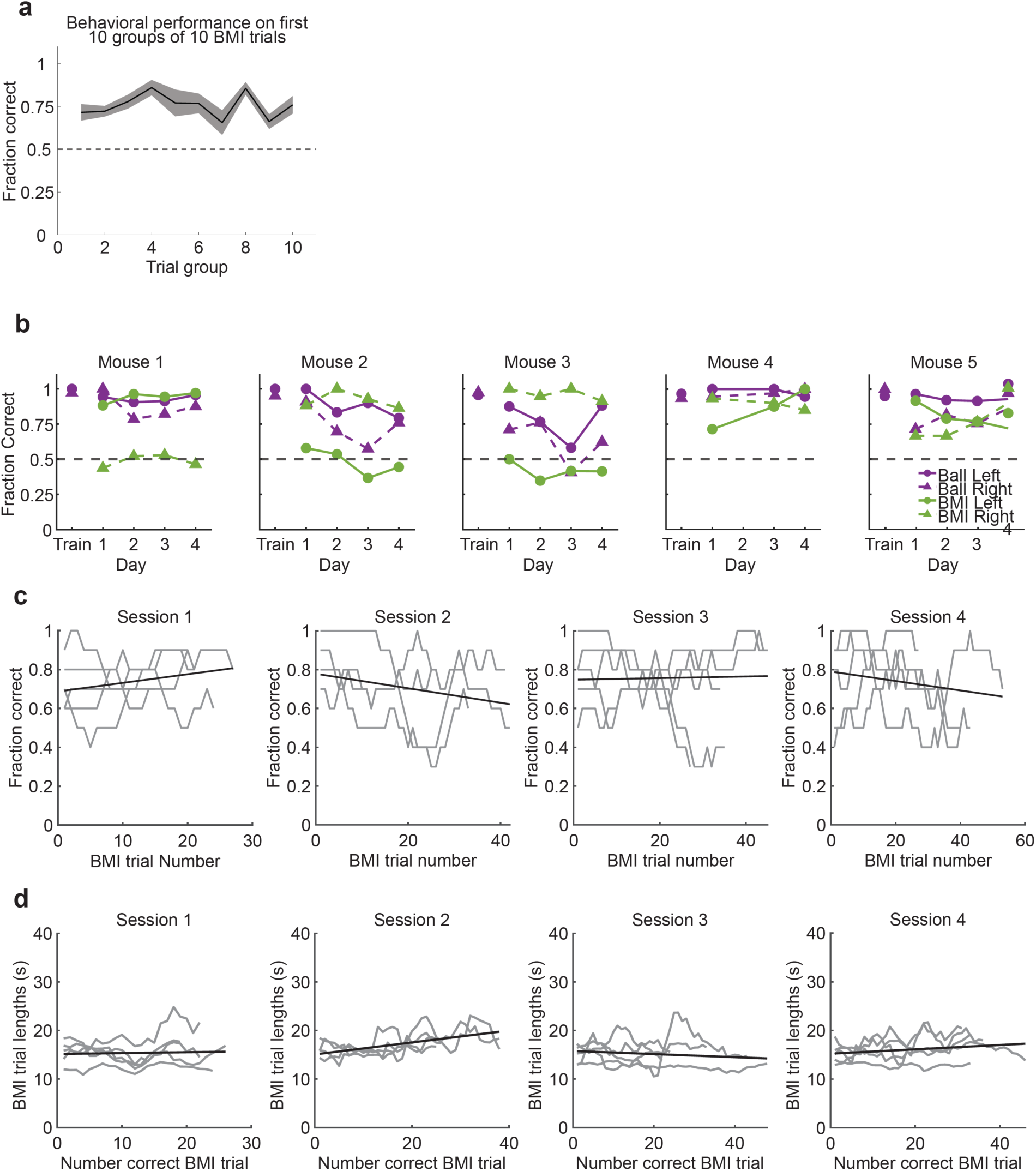
Detailed characterization of BMI performance over time. **a)** Performance of mice on the first 100 BMI trials, grouped into sets of 10 BMI trials. The mean ± SEM across sessions are plotted. **b)** Performance of mice plotted across sessions separately for each trial type and mouse. Each line corresponds to performance on a single trial type. **c)** Fraction correct during each session, smoothed with 10-trial sliding window; gray lines correspond to single animals and black line is line of best fit. **d)** BMI trial lengths during each session, smoothed with 5-trial sliding window; gray lines correspond to single animals and black line is line of best fit.

**Figure S3:**
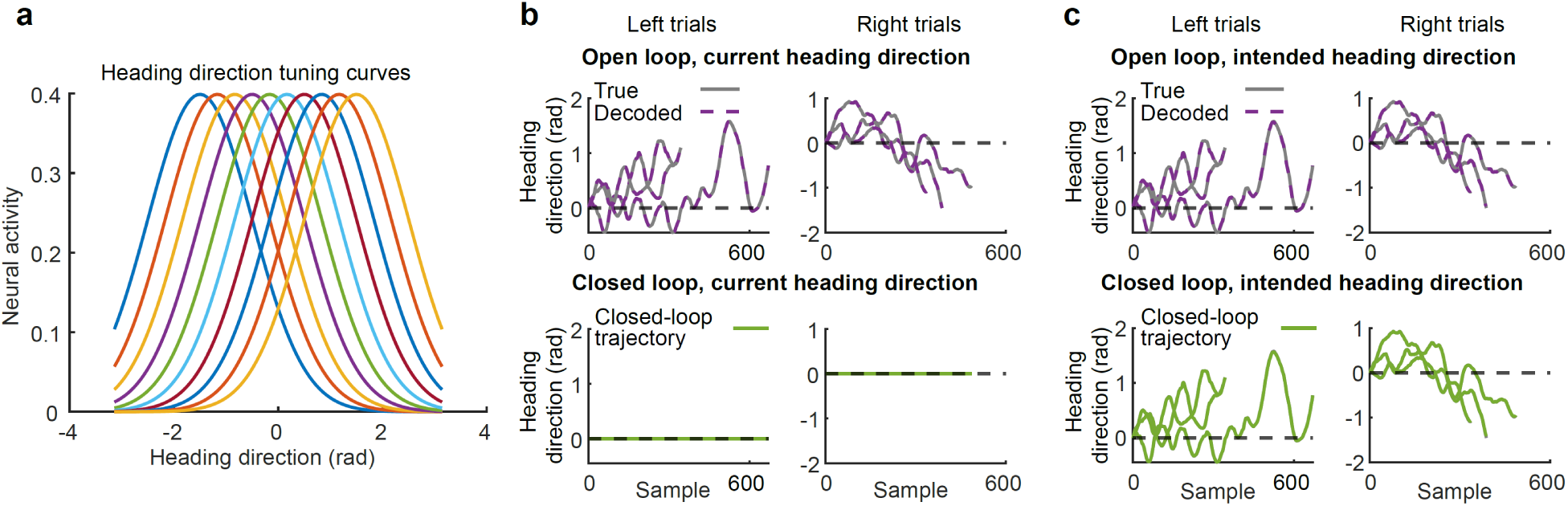
Modeling of task performance using different representations of heading direction. **a)** Model units had Gaussian tuning for heading direction; **b**) Top:In the open loop, when model units represent current heading direction, the agent navigates the maze and current heading direction can be accurately reconstructed from model unit activity. Bottom: In the closed loop, with model units representing current heading direction, the agent fails to turn in the maze; **c)** when model units represent intended heading direction at the subsequent time-step, open loop decoding accuracy remains high (**top**) but the agent can also update its heading direction and successfully complete trials in the closed loop (**bottom**)

**Figure S4:**
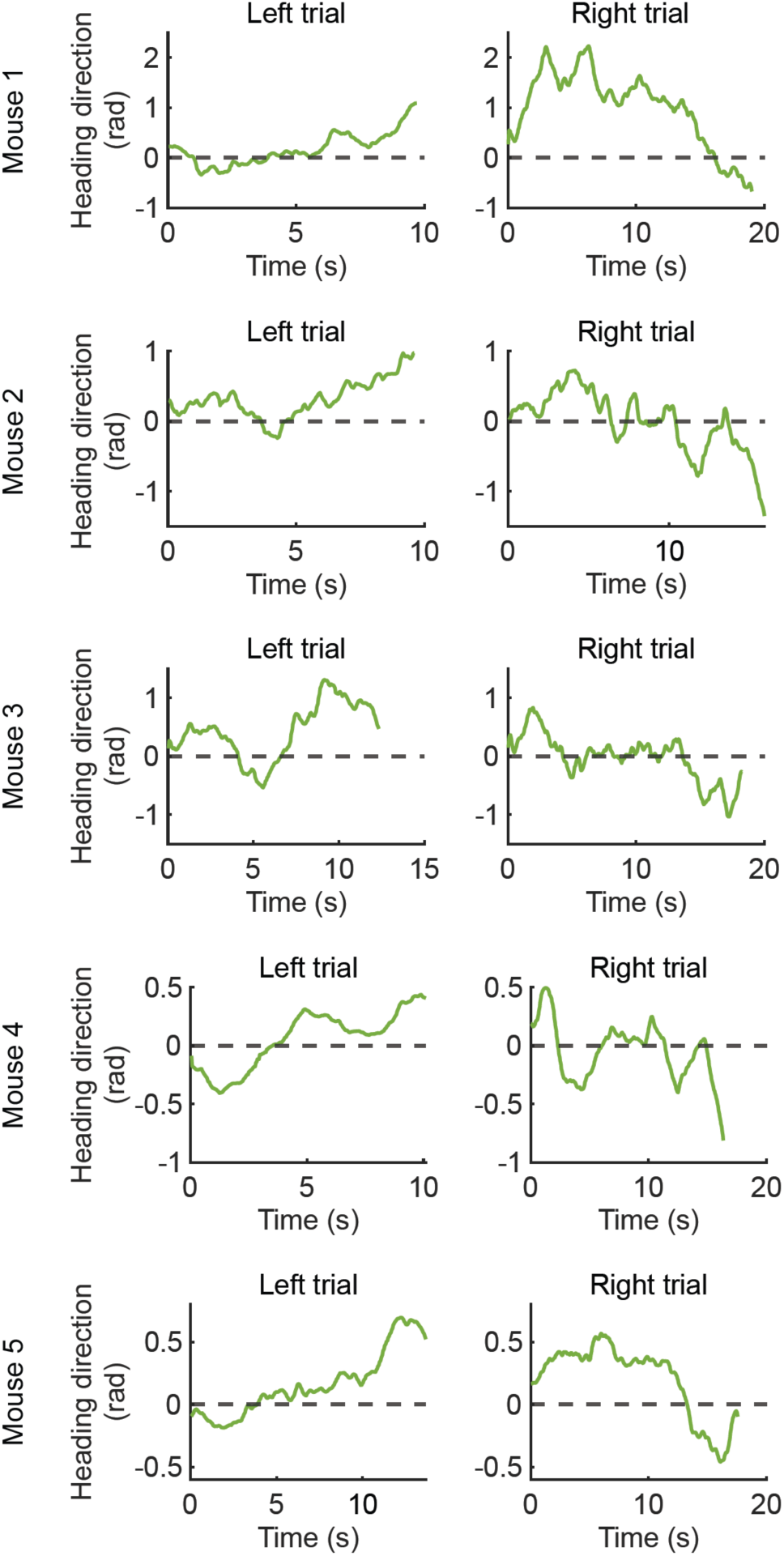
Examples of course corrections during BMI trials. Single trial examples of each mouse in the study making course corrections using the BMI.

**Figure S5:**
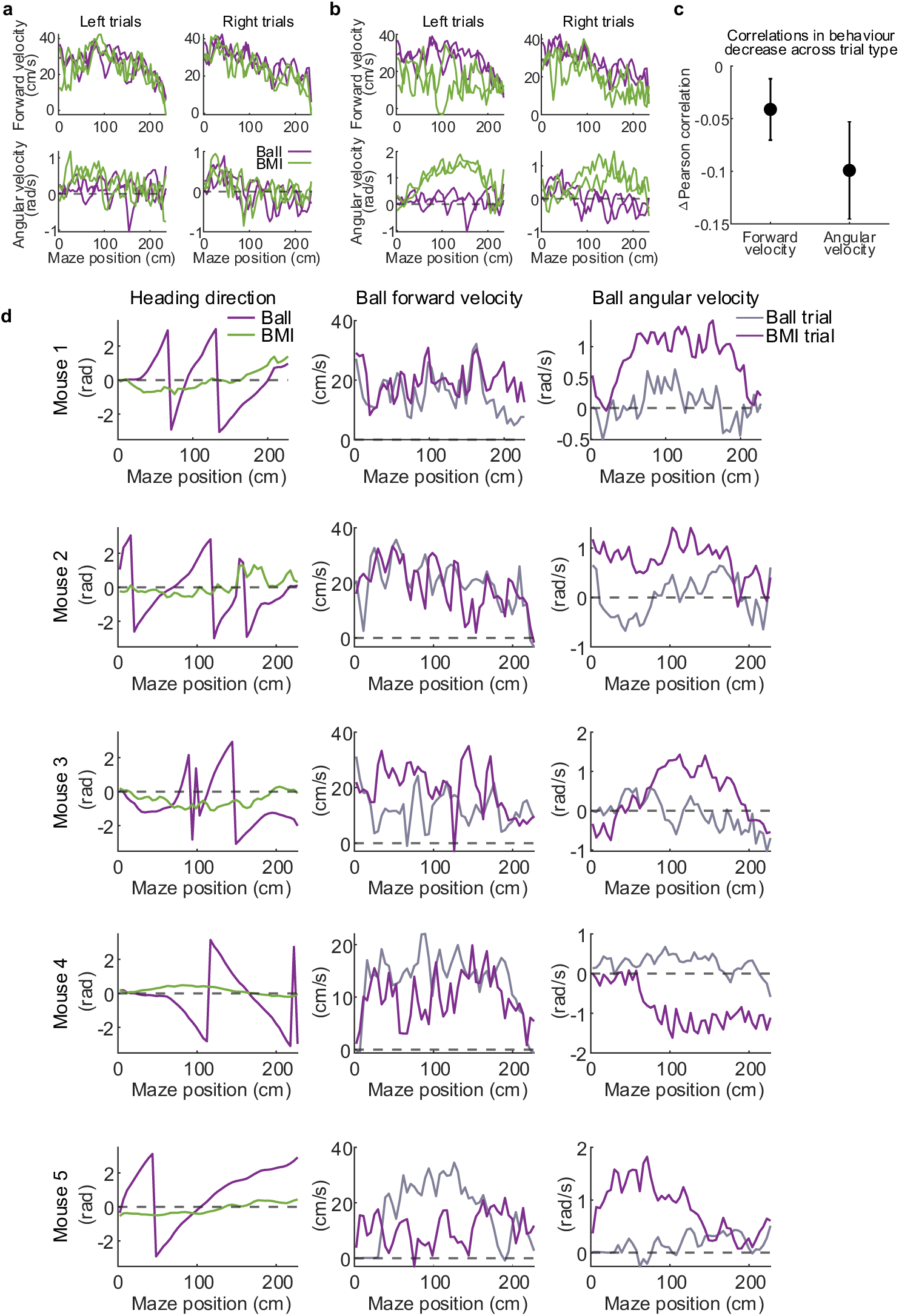
Characterization of differences in running patterns across Ball and BMI trials. **a)** Example spatially binned ball velocity trajectories of individual trials for mouse 1 session 1, for both ball and BMI trials. Examples of BMI trials that were similar to the selected ball trials are shown. **b)** Example spatially binned ball velocity trajectories of individual trials for mouse 1 session 1, for both ball and BMI trials. Examples of BMI trials that were different to the selected ball trials are shown. **c)** Differences between the correlations within and across trial types for forward and angular velocities. Data are plotted as mean ± 1 standard deviation across sessions. **d) Left column:** Example BMI trials where mice spin the ball along the yaw axis while traversing the maze. Green line indicates heading direction as controlled by decoder and magenta line indicates heading direction had it been controlled by the spherical treadmill. **Middle column:** Magenta line indicates example ball forward velocity during an example BMI trial where mice repetitively spun the ball along the yaw axis. Gray line indicates ball forward velocity in a typical ball trial. **Right column**: same as **middle column**, but for ball angular velocity.

**Figure S6:**
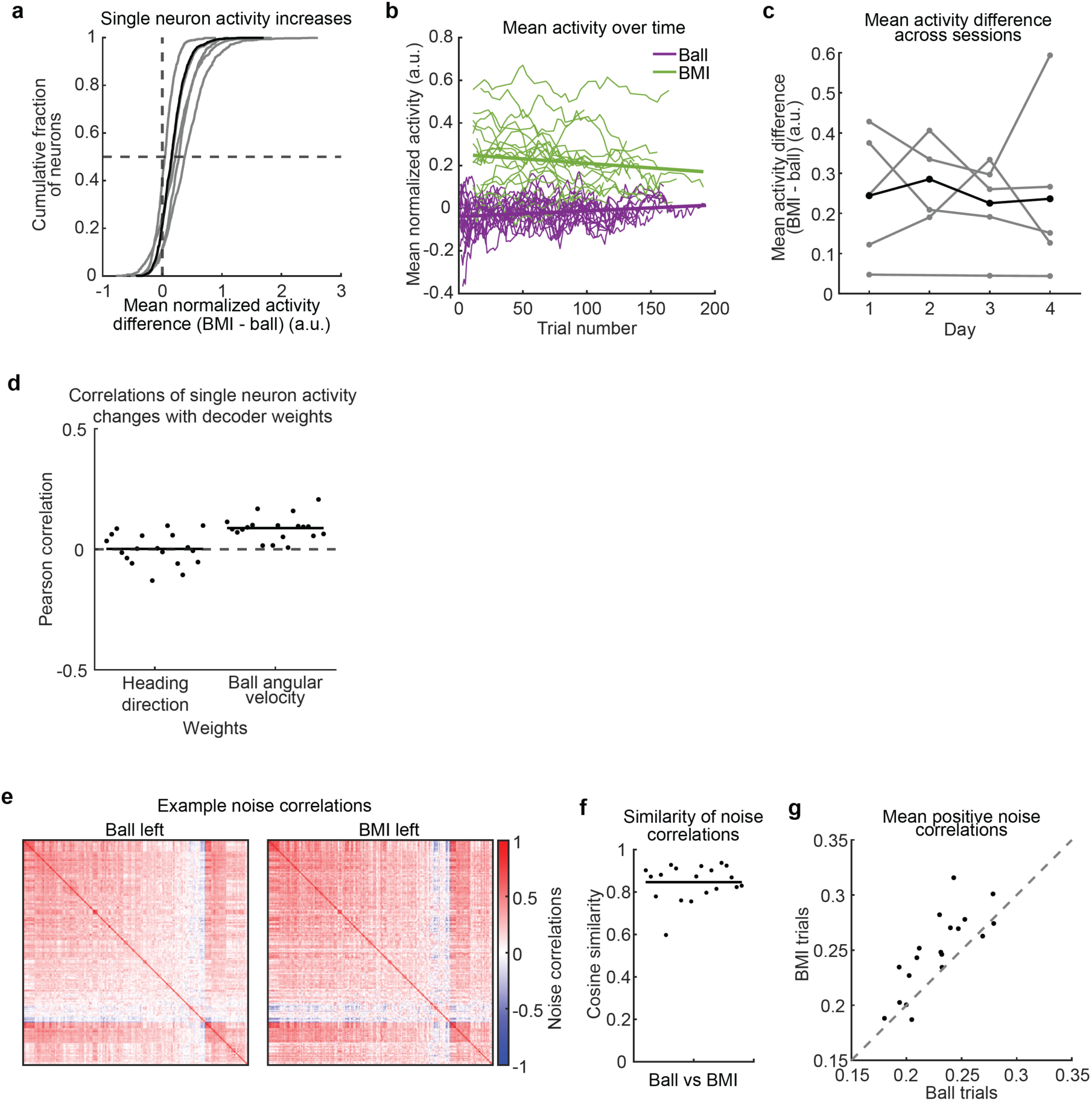
Differences in population activity between Ball and BMI trials. **a)** Distributions of change in mean normalized activity for individual neurons from ball to BMI trials. Grey lines are the means across sessions for individual mice, the black line is the mean over all mice. **b)** Mean normalized population activity plotted against trial number within sessions for ball and BMI trials. The thin lines are individual sessions, smoothed using a moving average across 10 trials. The thick lines are best fit lines. **c)** Mean population activity difference (BMI – ball) plotted against day of recording. Grey lines are the means across sessions for individual mice, the black line is the mean across mice. **d)** Correlations between the weights within neurons in view angle or ball angular velocity decoders with the absolute change in mean neural activity for individual neurons. Points are individual sessions, the black lines are means across sessions. **e)** Noise correlations between pairs of neurons on ball or BMI left trials for an example session. Neurons were clustered and sorted using the correlations for ball left trials, and the same sorting was used for both images. **f)** Cosine similarities between noise correlations on ball and BMI trials. Correlations and similarities were computed for left and right trials separately then averaged for plotting. Each point is an individual session, the line is the mean across sessions. **g)** Mean of neuron pairs with positive noise correlations on ball or BMI trials. Each point is the mean for an individual session.

**Figure S7:**
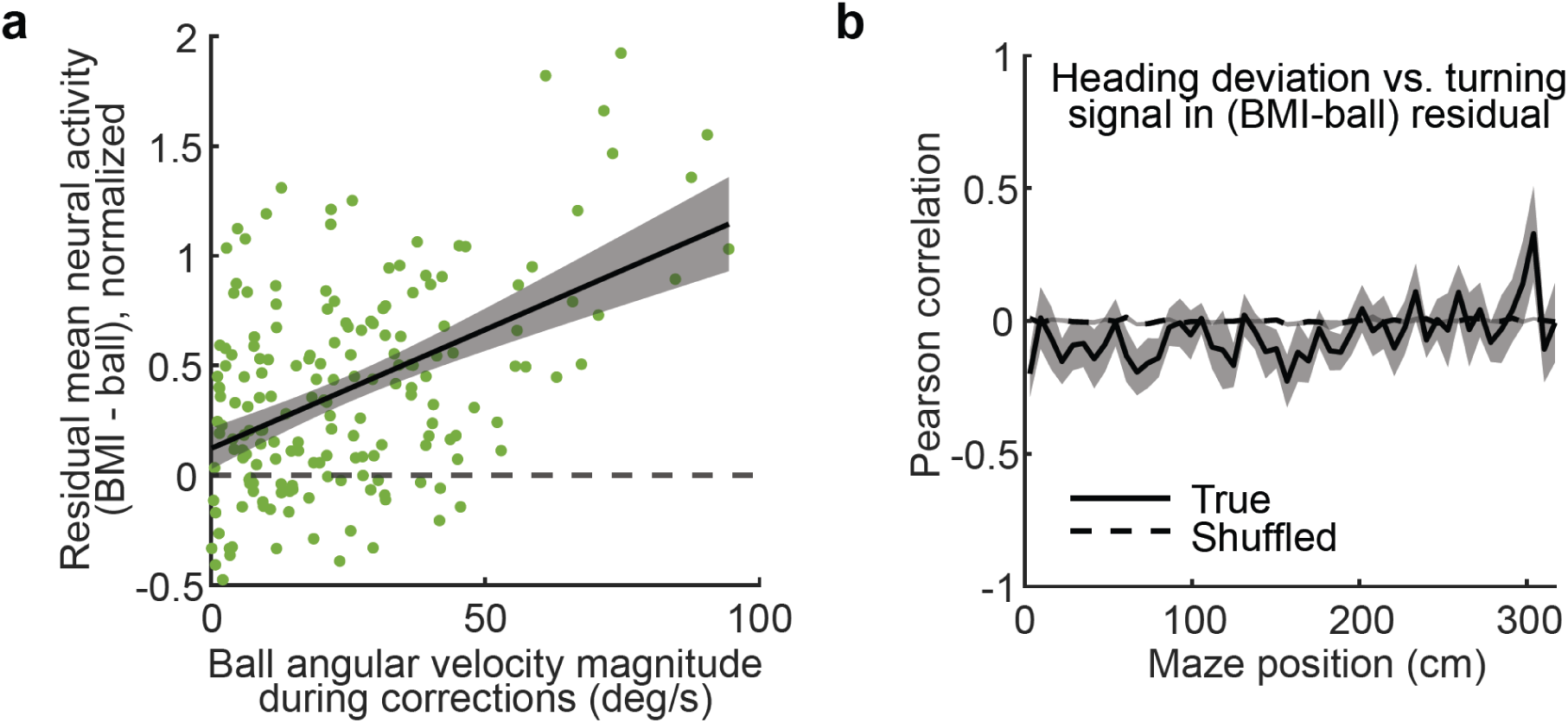
Residual neural activity during heading deviation corrections on BMI trials. **a**) Residual (BMI – ball) activity in spatial bins where motor corrective activity was detected, and normalized heading deviations were greater than 1, plotted against magnitude of angular velocity on ball. **b**) Correlation between heading deviation and decoded angular velocity, with the latter computed from decoded residual (BMI – ball) activity using the online BMI view angle decoder. Hierarchical bootstrap means and standard deviations across all sessions are shown for the true data (solid line) and for data with shuffled trial type labels (dashed line).

**Figure S8:**
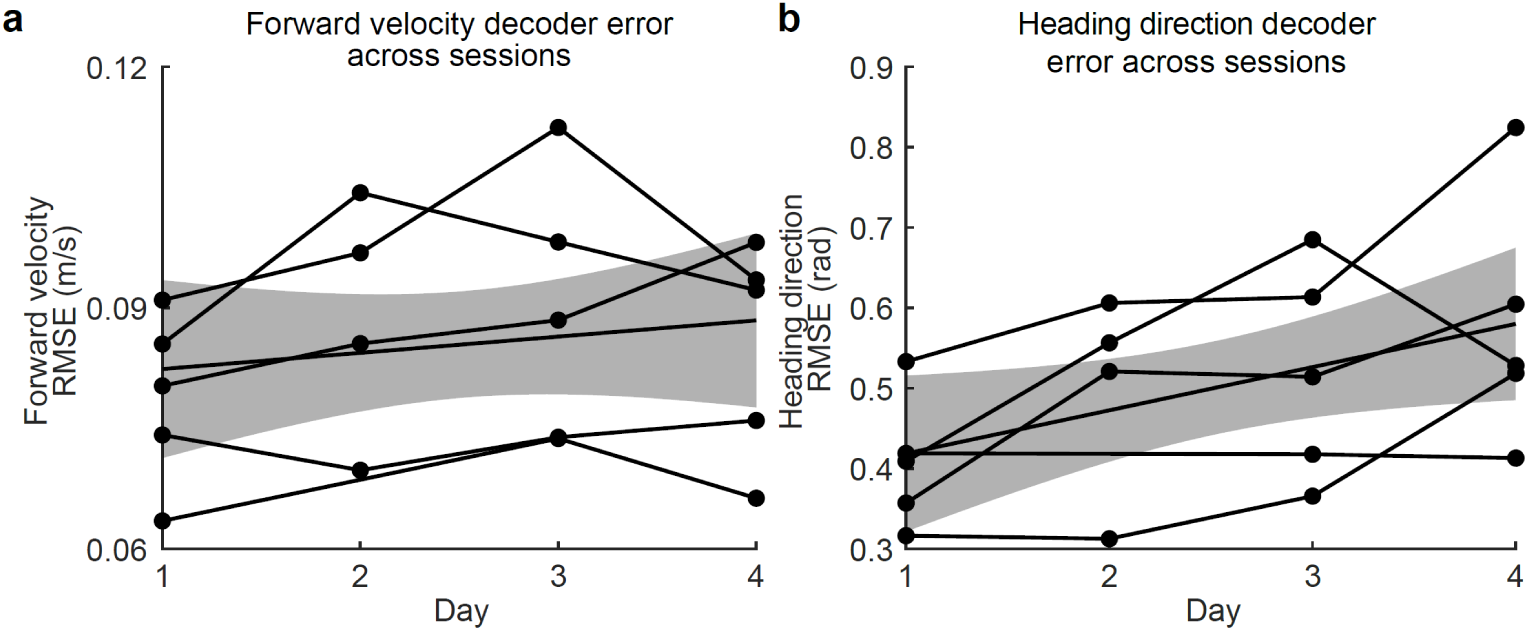
Decoder error accumulates across days. **a**) Forward velocity decoder accuracy measured using RMSE during ball trials across the four BMI testing sessions. Each line corresponds to a single mouse. A regression line with 95% confidence intervals is also shown. **b**) Heading direction decoder accuracy measured using RMSE during ball trials across the four BMI testing sessions. Each line corresponds to a single mouse; the shaded area corresponds to 95% confidence intervals of a linear regression.

## Methods

### Key resources table

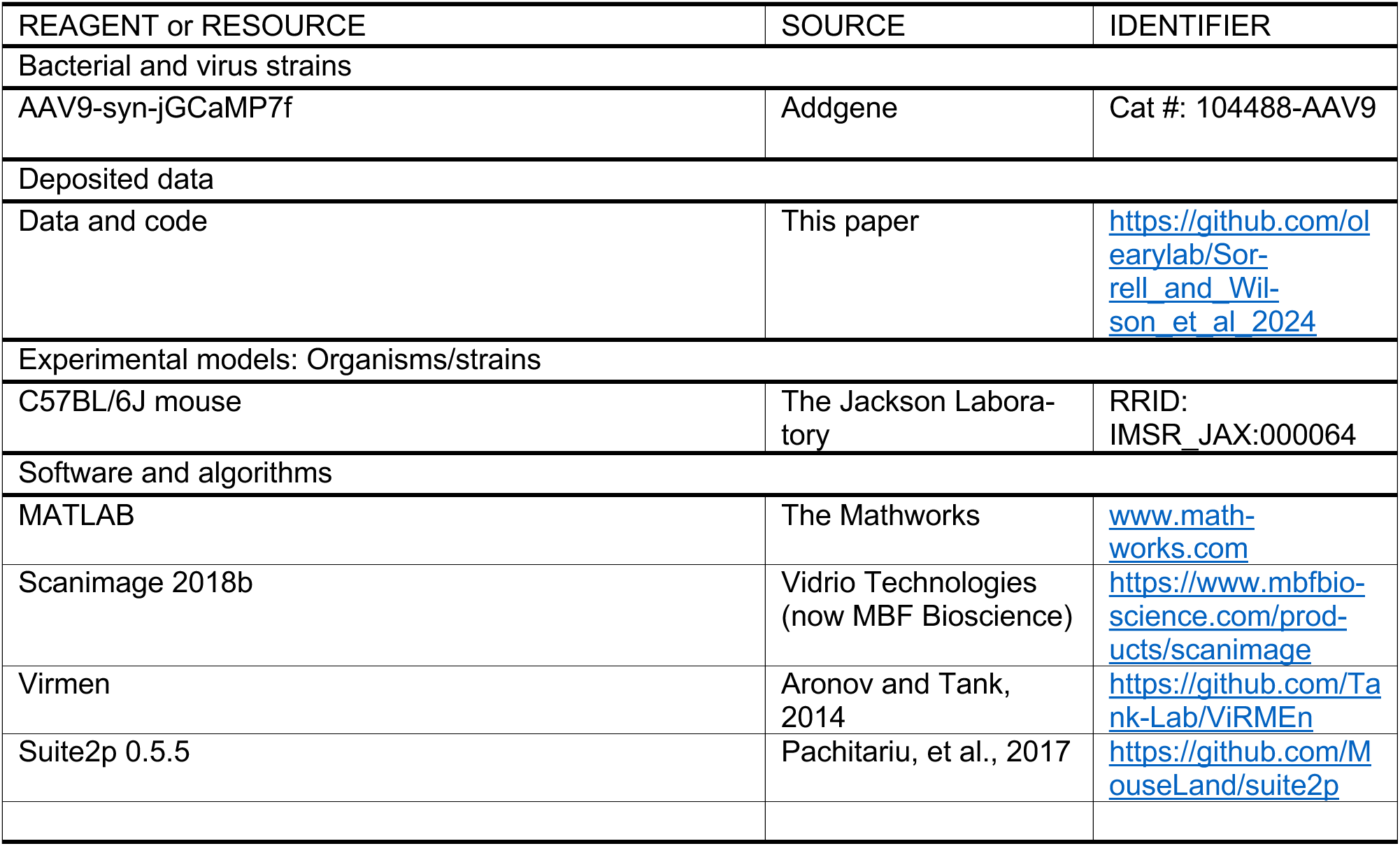

## Resource Availability

### Lead contact

Further information and requests for resources and reagents should be directed to and will be fulfilled by the lead contact, Timothy O’Leary.

### Materials availability

This study did not generate new unique reagents.

### Data and code availability

Data will be deposited and made available by the date of publication. Code used to obtain the results will be available at GitHub (https://github.com/olearylab/Sorrell_and_Wilson_et_al_2024) on the date of publication.

### Animals

All procedures involving animals were approved by the Harvard Medical School Institutional Animal Care and Use Committee (Protocol IS00000073-6). Behavior and imaging data were obtained from five male mice aged 8–12 weeks before the initial window/ virus injection procedure. Mice were obtained from The Jackson Laboratory (C57/BL6J, stock no.000664). Mice were group-housed when possible. The colony was maintained on a reversed 12-12 h light/dark cycle.

### Cranial window installation

Mice were administered dexamethasone (IP, 2 mg/kg) one to four hours prior to surgery. Anesthesia was induced using 3% isoflurane, and the mice were mounted in a stereotaxic head holder. A custom titanium headplate was installed over PPC and adhered to the skull with a 4:1 mixture of Metabond (Parkell) and carbon (Sigma Aldrich). A 3.2 mm diameter craniotomy was made centered on stereotaxic coordinates (−1.6 M/L, −2.2 A/P).

Virus was injected through pipettes (1.5mm O.D., 0.86mm I.D., GC150F-10, Harvard Apparatus) beveled at an angle of ∼30° with an outer diameter of 15 μm. AAV9-hSyn-jGCaMP7f (Addgene, diluted to 1–4 × 10^12^ GC/mL) was combined with Fast Green FCF (0.5 % w/v in PBS) at 9:1. The viral mixture was backloaded into the injection pipette and injected using air pressure. Injections were performed in a 3 × 3 grid with 400 μm inter-site spacing. The grid was centered at the coordinates −1.6 M/L, −2.2 A/P. At each injection site, the pipette was lowered to 250 μm below the pia and left in place for 1 min. 100 nL of virus was injected over ∼2 minutes, and the pipette was left in place for 2 minutes after the injection.

After the virus injections, the dura was removed from the region of the brain contained in the cranial window and a glass plug installed. The glass plug consisted of one 4 mm coverglass and two 3 mm pieces of coverglass (#1 Thickness, Warner Instruments). The coverglasses were adhered to one another by optical adhesive (NOA 81, Norland Products). The glass plug was glued in place using Insta-Cure (Bob Smith Industries) and then cemented with the Metabond/carbon mixture. We then cemented a metal ring to the headplate for light blocking and maintaining continuous immersion of the microscope objective.

### Two-Photon Imaging

The field of view was selected to contain PPC (centered at −1.7 M/L and −2.0 A/P) and be devoid of major blood vessels. Images were collected using a custom two-photon microscope. A Chameleon Vision II laser (Coherent) provided dispersion-compensated 920 nm excitation. The scan path used a resonant scanner and galvanometric mirror to perform bidirectional, full-frame scanning at ∼30 Hz. Power at the sample was always less than 50 mW as measured during scanning with flyback blanking enabled. The microscope was controlled by Scanimage 2018b (Vidrio Technologies).

### Virtual-Reality Behavior Setup

Full details for the virtual-reality behavior rig can be found at https://github.com/HarveyLab/mouseVR. In brief, the virtual reality environment was created, controlled, and displayed by ViRMEN (virtual reality mouse engine) software^46^. A laser projector (picobit, Celluon) displayed an image onto translucent plastic film. Mice moved through the virtual environment by rotating an 8” spherical treadmill made of expanded polystyrene. Optical sensors measured ball rotations along the pitch and yaw axes, and these signals controlled forward and angular velocity in the virtual world, respectively. Reward (6 g/L acesulfame potassium in tap water) was controlled by a solenoid circuit and delivered by lick spout.

### Behavioral Training

Mice underwent water restriction for at least 5 days before starting behavioral training. Mice were trained daily for approximately 60 minutes. The first three maze groups (A-C) consisted of T-Mazes in which the mouse needed to enter the T-arm instructed by the tower location in order to receive reward. In these mazes, mice became comfortable being on the ball and navigating by moving the ball. They also became familiar with the general structure of the maze (a long corridor followed by a T-junction and two opposing arms (**Figure 1a**; see **Figure 3a** for physical dimensions). The fourth maze group contained two sets of mazes: those from maze group C and those from maze group E. Maze group E is identical to maze group C, except that both arms of the T contain a tower so the mouse must turn based on the cue present in the corridor. In maze D, trial types can be from either maze group. Maze group F introduces a delay by replacing part of the cue texture on the corridor wall with a trial-type neutral texture. The delay is gradually lengthened over the course of days until it reaches the final length of 1/3 the T-corridor. Mice were previously used photostimulation experiments not described in this paper.

### Image Preprocessing

#### Downsampling

We first downsampled images by a factor of 4 (from 512 × 512 pixels to 128 × 128) with bicubic interpolation. This provided 4.8 μm/pixel resolution, with the diameter of putative cell bodies spanning 2–5 pixels (**Figure 1k–n**).

#### Image Registration

The precise anatomical location of the field-of-view varied, due to the mouse’s movements and minor mechanical drift within sessions and across days. To attenuate this variation, we used phase correlation to register each image frame to a fixed template (limiting transformations to translations). This template was constructed as the average fluorescence signal for each pixel, across all frames in the training dataset.

#### Temporal Filtering

We temporally filtered the images using an online ΔF/F filter. This filter fails when there are negative values (which may arise due to baseline subtraction during imaging). We therefore added a constant offset to all pixel values to guarantee positivity. The offset used was mouse specific. We performed this filtering using:

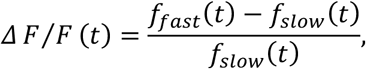

where *f*_fast_(*t*) and *f*_slow_(*t*) were the results of filtering the downsampled pixel values using a discrete-time exponential moving-average filter

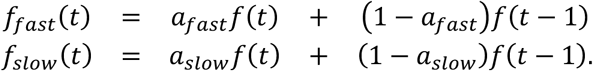

We determined the constants *a*_slow_ and *a*_fast_ from the time constants *τ*_fast_ = 0.5 s and *τ*_slow_ = 45 s, respectively, where *a* = 1 − *exp*(−1⁄*τ f_s_*), with the sampling frequency of *f*_s_ = 30 Hz. We chose these time constants to filter out low frequency changes, such as slow baseline shifts, as well as high frequency noise, while retaining useful calcium signals. This filter normalized and bandpass filtered the pixel fluorescence values. When run online, the moving-average ΔF/F filter required a burn-in period to provide accurate initial conditions for *f*_fast_ and *f*_slow_. For offline training, we achieved this by first filtering the data backwards. For online processing, we briefly ran the filter on some initial images in before starting the testing session.

#### Spatial Filtering

Following temporal filtering, we applied a spatial bandpass filter to the data. We subtracted a local background average, obtained using a Gaussian blur (radius *σ*_large_ = 5 downsampled pixels), from a local average (radius *σ*_small_ = 0.6 downsampled pixels).

### Training the Decoder

For the first cohort of mice (M1–3), mice performed trials normally using ball control for approximately 80 trials at the start of the training session. The mice then continued to run ball trials while the decoder was trained. We pre-processed fluorescence images as described above and appended a constant feature (set to 1) to each pixel vector. We also created a vector that indicated whether each data point was valid (to be used in training) or invalid (ignored for training). Invalid data included data from inter-trial intervals and incorrect trials. We then trained the decoder using the least mean squares (LMS) algorithm on the valid training set for 15 complete iterations through the set. We shuffled each matching pair of training samples in time on each iteration. We used the LMS update step:

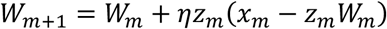

Where *η* was the learning rate, set to 10^−3^ (chosen empirically), *W* was the weights, *x* was the true kinematic variable, *z* was the processed neural activity vector, and *m* was the sample index. We initialized the weights to a set of weights from a previous days trained decoder.

For the second cohort of mice (M4, M5), we made some minor changes in the protocol. We balanced the number of left and right trials used for decoder training. Second, we set data during a trial where the mouse made more than a complete 360 degree rotation as invalid (as this was deemed poor behavior). Third, we initialized the weights to zero, with the bias term initialized to the mean of the variable being decoded. Finally, we only used five complete iterations to train the decoder because this was sufficient for convergence and reduced training time. In the implementation for the first cohort (subjects M1–3), we applied a fixed negative bias to the forward velocity used to train the BMI, equivalent to 0.01145 m/s, which is less than 10% the typical ball-controlled forward velocity (ranging from −0.23–0.63 m/s, with median forward speeds for each session spanning 0.11–0.20 m/s; c.f. **Figure 1e**). This bias had no substantive effect and was not included for the second cohort (M4, M5).

### Open-Loop Analysis

For assessing open-loop decoder performance in **Figure 1**, we used several days for each mouse where they performed only ball trials. For each day, we split the session into a training and testing dataset. We chose the training data set to be from the start of the session until the end of 40 correct left trials and 40 correct right trials. We used the rest of the session as test data. We then trained a decoder on the training data using the method for the second cohort of mice, as described above, and tested on the testing data by multiplying the processed testing image vector by the weight vector at each time point. We calculated the coefficient of determination, *R*^2^:

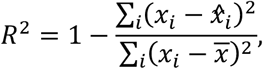

where *x* was the current true value, *x̂*_*i*_, was the current decoded value, and *x̄* was the mean of the testing dataset. We repeated this five times for each day, and calculated averages of these repeats.

For assessing performance using single neurons, we followed the same procedure as above but trained the decoder on the traces for ROIs detected using *Suite2p* that were identified as neurons by our CNN (**Methods:** *ROI segmentation*). The learning rate used was *η* = 10^-2^, and down-sampling, image registration and spatial filtering were not required.

### BMI Testing

On the day that we trained the decoder, we used the rest of the session for testing. In the testing section, the first ten trials were ball trials, to ensure the mouse was still engaged with the task. The eleventh trial was a BMI trial, and subsequently every third trial was a BMI trial, with the intermediary trials being ball trials. On subsequent days, we used this same testing protocol, but used the whole session for testing, as we used the same decoder across all days without any retraining. For assessing closed-loop performance, each mouse had one day of training and testing, followed by three days of testing using the same decoder trained on the first day.

During closed-loop decoding, we low-pass filtered the decoded heading direction using a first order FIR filter, with a constant *α* = 0.2:

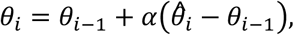

where *θ* is the filtered heading direction and 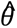 is the decoder output. We implemented a user function in Scanimage that wrote each decoder output to an analog output channel each time the microscope acquired an imaging frame. We closed the loop by routing these outputs to the data acquisition system controlling ViRMEN.

Different mice had different histories of BMI experience by the time the four-day stretch was recorded. We only report the results for the four days of a fixed decoder as this was consistently recorded across all mice. Also, we excluded the results for day 2 for M4 as the reward spout was not in place during this session.

## Data Analysis

### ROI segmentation

We used a previously published 3-layer Convolutional Neural Network (CNN)^26^ to classify sources that were automatically generated by *Suite2p*. Inputs to the CNN were 25-pixel windows surrounding each ROI mask, and outputs were labels indicating whether the source was a cell body with an acceptable shape.

### Spatial Binning

For much of the analysis and plotting, we binned trial data according to the linearized position of the mouse in the maze. We use 50 spatial bins, and excluded samples in inter-trial intervals. Linearized position was defined as using:

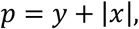

where *y* is the mouse’s forward position in the maze (along the T stem), and *x* is the mouse’s horizontal displacement within the left/right T-arms. For each trial, we averaged binned quantities over all samples where the mouse was located in a given spatial bin.

### Correlations of Behavioral Trajectories

In **Figure S3c**, we summarized within-condition (e.g. Ball-left, BMI-right) correlations by averaging the Pearson correlation between binned velocity (forward or angular) for all pairs of trials within each condition. We similarly assessed cross-control-type correlations as the average correlation between all trial pairs with the same turn direction, but different control types (Ball vs. BMI). We report “ΔPearson correlation” as the cross-control-type correlation, minus the within-control type correlation.

### Support-Vector Machine (SVM) Analyses

For **Figure 3f** and **Figure 6a**, we used the MATLAB function *fitclinear* to train SVMs for classifying trials as Ball or BMI. We trained a separate SVM for each location bin and trained a separate set of classifiers for left and right trials. For each session, we matched the number of training trials for each condition by sub-sampling each trial group to the lowest common number of trials for all trial types. We ran this analysis for 100 different sub-samples and report the average across all sub-samples. We assessed performance under 5-fold cross-validation (splitting trials in each session into five groups, and predicted trial type for each group using an SVM trained on the remaining four). For classifying trial type based on behavior (**Figure 3f**), we used the Z-scored ball forward and angular velocities. For classifying trial type based on neural data (**Figure 6a**), we used ΔF/F filtered and Z-scored fluorescence traces from identified neurons. We calculated the null distribution of SVM performance by shuffling the trial labels (100 shuffles). In all cases, we generated confidence intervals in cross-validated classification accuracy using a bootstrap procedure (**Methods:** *Hierarchical bootstrapping*).

### Heading Deviations

We used the mouse’s typical behavior during correct Ball trials from BMI training sessions to define a reference heading, which we interpret as the mouse’s intention. Within each session, we spatially binned heading (50 bins) and calculated the mean *μ*_*θ*,*i*_ and standard deviation *σ*_*θ*,*i*_ for each bin *i* over all correct Ball trials. For each BMI trial, we defined the per-bin heading deviation as *Δθ_i_* = (*θ_i_* − *μ*_*θ*,*i*_)/ *σ*_*θ*,*i*_, where *θ_i_* is the heading measured in the *i^th^* bin.

### Decoder Angular Velocities

We converted decoded heading direction to a heading-velocity command for the ViRMEN system by taking the discrete derivative 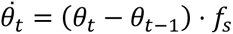, where *f*_s_ = 30 Hz is the sampling rate (the final angular velocity for each session was set to 0). We also used the decoded velocities to estimate angular velocities from decoder output during offline analysis.

### Correlations with Heading Deviations

In **Figure 4c,d**, we correlated the heading deviation with the angular velocity measured by the ball or BMI decoder. We only included samples in which the standardized heading deviation was large (|*Δθ*| > 1) and calculated correlation coefficients for each spatial bin separately.

### Neural Activity Normalization

For some offline analyses (**Figures 5, 6**) of isolated ROIs (putative neuronal cell bodies), we standardized the filtered ΔF/F fluorescence traces. We z-scored the ΔF/F time series for each neuron using the mean and standard deviation during correct ball trials only. Note that this normalization was not applied to signals used for BMI decoding. For **Figure 6b**, we used a qualitative normalization that scaled each mean tuning curve to the range [0,1] using the maximum (*c*_max_) and minimum (*c*_min_) values across both the ball and BMI tuning curves, with normalized amplitudes for the *i^th^* bin given as *c_norm_*_;*i*_, = (*c_max_* – *c_i_*)/(*c_max_* − *c_min_*).

### Correlations of Neural Activity with Running Movements

In **Figure 5a,b**, we correlated neural activity with running movements during periods with heading deviations. We binned normalized neuronal activity, heading deviations, and ball angular velocity into 50 spatial bins. We then correlated the neural activity (either individual neuron, or the mean across the population of neurons) with the ball angular velocity. We restricted this analysis to samples in which the magnitude of the normalized heading deviation was greater than one, |Δ*θ*| > 1, and in which the ball angular velocity had the opposite sign to the heading deviation, i.e. the behavior was an attempt to correct the heading deviation. We discarded any bins that were within five bins of another accepted bin in order to reduce the impact of the temporal dependence of samples. We calculated correlations separately for each session and also calculated correlations when shuffling neural data relative to behavioral data to compute the significance of the true correlations.

### Angular Velocity Decoding

We trained pixel-based decoders to decode ball angular velocity during corrective movements in correct BMI trials. We defined corrective movements as times in which the ball angular velocity was in the opposite direction to the heading deviation. For this analysis, we defined the standardized heading deviation for each time-point *Δθ*(*t*) using the mean and standard deviation for the spatial bin occupied at time *t* (**Methods:** *Heading Deviations*).

In **Figure 5c**, we assessed decoder accuracy separately for samples in which heading deviations were small |Δ*θ*| ≤ 1 and large |Δ*θ*| > 1. We restricted the analysis to correct BMI trials and used the same training methods described in **Methods:** *Training the Decoder*, but with 20 complete iterations through the training data. We used five-fold cross-validation and sub-sampled the data trial-wise (without replacement) to ensure that all sessions and left-vs. right-turn conditions had an equal number of training and testing trials. To account for variability in this sub-sampling, we repeated the analysis for 30 different sub-samples. We report the average decoder accuracy for the testing group for each set of weights over all samples.

### Bootstrapping

When plotting neural tuning curves (**Figure 6d**) and behavioral trajectories (**Figure 3c,e**), we used a bootstrap to estimate confidence intervals. We binned each trial into 50 spatial bins. We set the number of trials (*n*) for each bootstrap resample to the minimum of the number of correct trials across all four (Ball/BMI, left/right) trial types. We drew 100 bootstrap samples of *n* trials (with replacement) from the pool of correct trials for each trial type and calculated the mean and 95% confidence intervals from these samples.

### Tuning Curves

When calculating neuronal tuning curves (**Figure 6b,d**), we first preprocessed and binned neural data for each trial into 50 spatial bins. We then calculated position tuning curves via bootstrap resampling, as described above.

### Mean-Activity Changes

To calculate changes in mean neural activity levels in **Figure 6c** and **Figure S4a**, we first preprocessed and normalized the neural data as above. We then calculated the mean activity during valid data samples (i.e. not in the inter-trial interval) of correct trials for each neuron for each trial type, and the mean of these across all neurons. Additionally, we spatially binned the neural data and calculated the bootstrapped resampled mean for each trial and neuron and averaged these across neurons and trials within each trial type to get a population average in each of the 50 spatial bins.

### Tuning Curve Changes

To compute tuning curve changes in **Figure 6d,e**, we first calculated positional tuning curves as in **Methods:** *Tuning Curves*. We used a shuffle test to determine which neurons were significantly tuned to at least one type of trial. We used a circular shuffle to assess the per-neuron noise level while preserving temporally correlated slow fluctuations at the trial and session timescales. We masked out BMI-control periods, and then sampled 1000 circular shuffles (with each sample shifting the data by at least 10 seconds). For each shuffle, we computed the average shuffled activity in each spatial bin (50 bins) during Ball trials. We used the maximum mean activity over all spatial bins for each shuffled sample and took the 99^th^ percentile of these maxima as the threshold for significant tuning. We deemed a neuron to have significant position tuning if, for a given trial type, the maximum value of its mean tuning curve was above this threshold.

In **Figure 6e**, we summarize the prevalence of various qualitative changes in neuronal tuning between Ball and BMI trials. If a neuron had significant tuning on BMI trials but not ball trials for at least one direction, this neuron was said to have gained tuning. If a neuron had significant tuning on ball trials but not BMI trials for at least one direction, this neuron was said to have lost tuning. If a neuron had significant peaks on both trial types for at least one direction, we then also assessed whether the tuning peak had a change in location, amplitude, or width. The location of the peak was the bin of the tuning curve with the maximum value, the amplitude was the maximum minus the minimum value of the tuning curve, and the width was the number of bins of the tuning curve that were above the significance threshold. These values were calculated for each bootstrap sample of the tuning curves for ball and BMI trials for each direction. A parameter was deemed to have changed if the mean of that parameter on ball trials was outside the 95% confidence interval around the mean for that parameter on BMI trials, and vice versa. If a neuron had significant peaks on both trial types for at least one direction, but no significant change in any of the parameters, this neuron was said to have no change in tuning. We computed the fraction of neurons separately for left and right turn trials, and then calculated the mean across the turn directions for each session.

### Correlations of weights and mean activity changes

For **Figure S4d,** we first calculated the mean activity change for each neuron. We did this by subtracting the mean activity during valid samples of correct ball trials from the mean on correct BMI trials, and taking the absolute value to obtain the magnitude of the change in mean activity. Then we correlated these changes with the magnitude of the weights for each neuron (calculated from the pixel decoders) for either the online view angle decoder, or the offline corrective angular velocity decoder. In the latter case, we calculated correlations for each cross-validation and subsample, before calculating the mean of the resulting correlations.

### Noise Correlations

In **Figure S4e-g**, we calculated noise correlations from preprocessed and normalized neuronal fluorescence traces. We computed the correlation between activity of each pair of neurons separately for each spatial bin in correct trials and then averaged across all bins. In **Figure S4f**, we computed cosine similarities of the matrix of noise correlations for Ball and BMI trials. We analyzed left and right trials separately and then averaged these two correlation coefficients into a single summary statistic. In **Figure S4g**, we restricted analysis to positive correlations, plotting the mean over only neuron pairs with positive correlations.

### Linear Models of Neural Activity

We trained linear regression models to predict the activity of individual neurons using behavioral variables during ball or BMI trials and tested the accuracy of the model predictions on the same or different trial types. We trained models separately for left and right trials. For each session, we matched the number of correct trials where mice did not perform any complete rotations (360 degree) for each trial type by sub-sampling each trial group to the lowest number for all trial types. We also removed all samples during the inter-trial interval from each trial. We ran this analysis for 30 different subsamples of trials and calculated the mean across subsamples.

We used the ball pitch, roll, and yaw velocities or 10 gaussian bumps created from the linearized position as the predictor variables. These position bumps were created by passing the linearized position through 10 different Gaussian pdfs, with means spaced evenly between the minimum and maximum values of linearized position, and standard deviations equal to half the distance between each mean. We then z-scored each predictor variable. We also appended a vector of ones to the predictor matrix to include a bias term.

We trained linear regression weights to predict ΔF/F filtered fluorescence traces of each neuron identified using Suite2p. We also either z-scored these traces across both ball and BMI trials for the current subsample and direction, or z-scored ball and BMI trials separately. When normalizing across both types, we maintain any differences in mean activity that exist across types. When normalizing within each type, we removed these mean shifts.

We trained the model weights using L2 regularized linear regression. We determined a suitable value of *λ* for each model by calculating the cross-validated accuracy on the same trial type for a single subsample of trials. We selected the value of *λ* that produced the lowest RMSE from a range of *λ* = [0.001,0.01,0.1,0,1,10,100,1000]. For each neuron and trial type/direction combination, we then computed the regularized regression weights using the chosen *λ*.

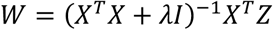

where W was the weight matrix, X was the matrix of training predictor variables, I was the identity matrix, and Z was the vector of neural activity for the current neuron. We performed 5-fold cross-validation and used the predictions of held-out data of each cross-validation as the within type test predictions. Predictions were made using matrix multiplication.

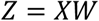

We computed the RMSE across all these predictions.

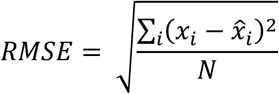

*x_i_* was the true value, and *x̂*_*i*_ the decoded value for sample i. N was the total number of testing samples. For across-type testing, we calculated the predictions for the entire test set with each cross-validated set of weights, then calculated the RMSE for each, and calculated the mean over the five models. We calculated the mean across sub-samples, directions and neurons for plotting in **Figure 6g-h**.

### Correlations of Residual Activity With Corrective Movements

In **Figure S7a,b**, we defined the “residual neural activity” as the difference between the neuronal activity during BMI-control and Ball-control trials, treating left- and right-turn trials separately. In **Figure S7a**, we used the preprocessed and normalized pixel fluorescence traces, and calculated the mean activity 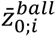 in 50 spatial bins (indexed by *i*) during correct Ball trials. We then subtracted this Ball-control reference from the binned activity for each BMI trial to generate residual neural activity 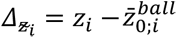 for each BMI trial. We then correlated location-binned residuals 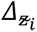 with ball angular velocities 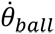 in bins where the ball angular velocities were corrective (opposite sign to heading deviation) and heading deviations were greater than 1, |Δ*θ*| > 1. We discarded any bins that were within five bins of another accepted bin in order to reduce the impact of the temporal dependence of samples.

### Correlations of Residual Projections and Heading Deviations

In **Figure S7b**, we defined the residual activity as 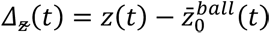, where 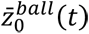 is the reference preprocessed neural activity for the location bin (without normalization) occupied by the mouse at time *t*. We passed 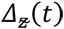 through the pixel-based BMI view-angle decoder, and differentiated to get the projection of residual activity onto turning (angular) velocity. We averaged these angular velocities within 50 spatial bins and correlated these binned velocities with heading deviations Δ*θ* separately for each bin. (only included samples where the magnitude of the normalized heading deviation was greater than one, |Δ*θ*| > 1).

### Hierarchical Bootstrapping

We used hierarchical bootstrapping^47^ for many plots, statistical calculations, and assessment of statistical significance. The hierarchical bootstrap allows us to estimate confidence intervals without specifying assumptions about the dependency between sessions from the same mouse or trials from the same session. For all tests, we drew 1000 bootstrap samples. We first sampled at the level of mice, then at the level of sessions (with replacement). We used the largest number of sessions across mice as the sample size for the lower level (this was four for both offline and online testing). Data are reported as mean ± standard error of the mean (SEM) of these hierarchical bootstrap distributions unless otherwise stated.

To test whether statistical quantities differed across conditions, we exploited that data were always paired. For example, if we observed that a summary statistic *ϕ* was larger in condition *B* compared to condition *A* trials, **ϕ*_B_* > **ϕ*_A_*, we performed a paired test by bootstrap sampling the difference *ϕ_A_* – *ϕ_B_* (rather than an unpaired test on independent samples from *ϕ_A_* and *ϕ_B_*). We made no *a priori* assumptions about the sign of any effect, interpreting all tests as two-tailed tests. For a significance threshold of *α*, we would reject the null if *p*_boot_ := *Pr*[*ϕ_A_* – *ϕ_B_* ≥ 0] < *½α*. In instances where *ϕ_A_* – *ϕ_B_* < 0 for all 1000 samples we report, *p*_boot_ < 0.001. When testing statistical significance for effects across 50 location bins, we applied the Benjamini-Hochberg^48^ correction procedure for a False-Discovery Rate (FDR) of α = 0.05 to the boostrapped two-tailed p-values.

## Author contributions

E.T.S., D.E.W., C.D.H., and T.O. conceived experiments. E.T.S., D.E.W. and M.E.R. built the BMI. D.E.W. and H.Y. conducted experiments. E.T.S., D.E.W., and M.E.R. analyzed the data with guidance from F.F., C.D.H., and T.O. E.T.S., D.E.W., C.D.H., and T.O. interpreted the results and wrote the paper with input from all authors.

## Acknowledgments

We thank members of the Harvey and O’Leary labs for useful discussions. AAV-syn-jGCaMP7f-WPRE was a gift from Douglas Kim & GENIE Project (Addgene viral prep # 104488-AAV9; http://n2t.net/addgene:104488; RRID:Addgene_104488). We thank the HMS Research Instrumentation Core for help with constructing behavior training rigs and Georg Jaindl for advice on real-time processing in Scanimage. This work was supported by NIH grants to D.E.W. (F32MH118698) and C.D.H. (DP1MH125776, R01NS089521), by the Harvard Mahoney Neuroscience Institute Fund (D.E.W.) and by a Human Frontier Science Program grant to C.D.H. and T.O. (RGY0069). E.T.S. was supported by funding from the Centre for Integrative Neuroscience Discovery (CIND), and the Cambridge Commonwealth, European & International Trust.

